# Multiplex design and discovery of proximity handles for programmable proteome editing

**DOI:** 10.1101/2025.10.13.681693

**Authors:** Chase C. Suiter, Green Ahn, Melodie Chiu, Yi Fu, Shayan Sadre, Jessica J. Simon, David S. Lee, Douglas M. Fowler, Dustin J. Maly, David Baker, Jay Shendure

## Abstract

Although we now have a rich toolset for genome editing, an equivalent framework for manipulating the proteome with a comparable flexibility and specificity remains elusive. A promising strategy for “proteome editing” is to use bifunctional molecules (*e.g.* PROteolysis-Targeting Chimeras or PROTACs^1^) that bring a target protein into proximity with a degradation or stabilization effector, but their broader application is constrained by a limited repertoire of well-characterized target or effector “handles”. We asked whether coupling *de novo* protein design to a multiplex screening framework could address this gap by accelerating the discovery of effector handles for intracellular protein degradation, stabilization, or relocalization. Using LABEL-seq^2^, a sequencing-based assay that enables multiplex, quantitative measurement of protein abundance, we screened 9,715 *de novo* designed candidate effector handles for their ability to recruit a target protein to components of the ubiquitin–proteasome system^3^ (UPS) (FBXL12, TRAF2, UCHL1, USP38) or the autophagy pathway^4^ (GABARAP, GABARAPL2, MAP1LC3A). In a single experiment, we discovered hundreds of *de novo* designed effector handles that reproducibly drove either intracellular degradation (n = 277) or stabilization (n = 204) of a reporter protein. Validation of a subset of these hits in an orthogonal assay confirmed that sequencing-based measurements from the primary screen reliably reflected changes in intracellular abundance of the target protein. Successful effector handles were discovered for both the UPS (n = 194) and autophagy (n = 287) pathways, which provide complementary routes for programmable proteome editing. Autophagy-recruiting effector handles generalized to endogenous targets, as substituting the reporter-specific target handle with a high-affinity MCL1 binder^5^ reduced endogenous levels of this intracellular oncoprotein^6^. Moreover, directing autophagy-recruiting effector handles to the outer mitochondrial membrane dramatically perturbed mitochondrial networks in a manner consistent with synthetic tethering and sequestration^7,8^. Beyond generating a diverse repertoire of protein abundance or localization effector handles, our results establish a scalable, low-cost platform that links deep learning–guided protein design to functional cellular readouts, and chart a course toward a general framework for programmable proteome editing.

## INTRODUCTION

The past decade has seen transformative advances in genome editing, with CRISPR-based technologies^9^ now enabling the precise, programmable manipulation of virtually any genomic locus (DNA) and, increasingly, transcribed nucleic acids (RNA). However, our ability to manipulate the remaining layer of the central dogma—proteins—with comparable flexibility remains primitive. Unlike nucleic acids, amino acid polymers lack predictable base-pairing rules, and thus resist the kinds of programmable targeting that have made genome and transcriptome editing routine. Advancing from genome editing to proteome editing will require not only a “phone book” for targeting proteins but also generalizable means of modulating their abundance, localization, and activity with precision within living cells and tissues.

One promising strategy for proteome editing is proximity induction^10^, in which a target protein is brought into physical proximity with a cellular effector to trigger a desired outcome—analogous to how guide RNAs and Cas9-effector fusions recruit chromatin-modifying activities to specific genomic loci. Targeted protein degradation with PROTACs^1^ exemplifies this concept^11^. By recruiting a protein-of-interest to endogenous degradation machinery, these bifunctional small molecules can selectively eliminate intracellular proteins. However, despite their therapeutic promise, most PROTACs rely on a narrow set of effector “handles” (e.g., CRBN, VHL), leaving the vast majority of the human E3 ligase repertoire—as well as other potential effector pathways—untapped. PROTACs’ broader application is further limited by the requirement for well-defined binding pockets on both the target and the effector, which are not always available, and by the challenge of scaling high-affinity ligand discovery across the proteome. As a result, the broader potential of proximity induction as a strategy for proteome editing is constrained not by concept, but by execution^12–14^.

Deep-learning–based *de novo* protein design^15,16^ offers a potentially powerful route to overcoming these constraints. Computational design tools can now rapidly generate candidate binders to virtually any surface epitope, bypassing the structural constraints that restrict small molecules and, in principle, enabling programmable recruitment of almost any target protein to a broad spectrum of effector functions defined by the nature and location of that recruitment. This potential is underscored by numerous examples of bifunctional proteins that use natural binding domains, antibody fragments, or combinations thereof for proximity induction^17–20^, yet these too remain limited by the range of available reagents. In principle, *de novo* protein design opens the door to the generalization of proximity induction-based proteome editing. However, while modern computational pipelines can now readily generate thousands of candidate binders against virtually any protein, most efforts to date have focused on screening for *in vitro* affinity rather than intracellular function.

A central challenge is how to scale functional screening to match the scale of *de novo* design. For example, if the goal is to build bifunctional proteome editors that modulate protein abundance, this requires quantitatively assessing which designs successfully recruit and degrade a target protein inside cells. Historically, methods for the multiplexed quantification of protein abundance have been limited. However, LABEL-seq^2^ offers a powerful solution. By linking the abundance of a target protein to an RNA barcode, LABEL-seq enables protein levels to be measured across thousands of cellular contexts within a single experiment.

Here we describe how integrating three key elements—*de novo* protein design, intracellular RNA barcoding of proteins, and the cell as a compartment in which designs and barcoded targets are brought together—creates a framework for scaling the discovery of reagents for intracellular proteome editing. We show that this combination enables sequencing-based quantification of how a target protein’s abundance is modulated by each of ∼10,000 candidate effector handles in one experiment, allowing not only detection of successful designs but also their ranking by intracellular potency, characterization of their effect-size distributions, and comparisons across target–effector pairings or cellular contexts. Together, these capabilities bridge protein design and cell-based functional screens, opening a path to the generation of a “phone book” for programmable proteome editing in human cells.

## RESULTS

### A strategy for the multiplex, intracellular assessment of *de novo* designed proteome editors

We set out to develop a multiplex workflow that quantitatively measures how libraries of bifunctional proteins—each comprising a target-binding domain (“target handle”) and an effector-binding domain (“effector handle”)—alter the intracellular abundance of a target protein. Classical proteomic methods (*e.g.* Western blots, mass spectrometry) are not readily scaled or multiplexed. Coupling fluorescence-activated cell sorting (FACS) with nucleic acid barcodes (“sort-seq”)^21,22^ provides an attractive alternative, but typically relies on a limited number of bins, reducing quantitative resolution, and is also constrained to a single fluorescently labeled target or effector per experiment.

To overcome these limitations, we turned to LABEL-seq^2^, which converts intracellular protein abundance into a sequencing-based readout. In this approach, a protein-of-interest (POI) is fused to an RNA-binding domain (RBD) and co-expressed with a uniquely barcoded RNA species that binds the RBD via an MS2 hairpin-MS2 coat protein (MCP) interaction^23^. Following immunoprecipitation (IP) of the RBD–POI fusion, the co-enriched RNA barcodes are sequenced, yielding a digital, quantitative measurement of protein abundance. In a *cis* configuration, sequence variants of the POI are tagged with unique barcodes, *e.g.* enabling multiplex measurement of their effects on protein stability^2^ (**Fig. S1A**). However, in a *trans* configuration, an invariant, RBD-tagged target can be labeled with an RNA barcode that instead encodes the identity of a concurrently delivered perturbation, *e.g.* a bifunctional proteome editor designed to modulate the RBD-tagged target’s abundance (**Fig. S1B**).

A critical feature of the *trans* configuration setup is that the cell itself serves as a compartment that links a specific perturbation (here, a bifunctional proteome editor) to a specific RNA barcode. As the RNA barcode remains bound to the target protein even after cells are lysed via the strong MS2-MCP interaction^2^, the intracellular effects of many candidate editors can potentially be measured in a single multiplex experiment in which the primary readout is massively parallel sequencing. Although this strategy is compatible with the screening of either target handles, effector handles, or combinations thereof, our proof-of-concept focused on evaluating computationally designed effector handles. In particular, we sought to evaluate thousands of designed binders to E3 ligases or autophagy-related proteins for their ability to mediate the intracellular degradation or stabilization of a target protein.

A first key component of our experimental setup for this proof-of-concept is a plasmid library encoding thousands of barcoded bifunctional proteome editors (**Fig. 1A**). In brief, a constant target handle (NbALFA, a high-affinity antibody to the ALFA tag^24^) is fused to a library of designed effector handles (**Fig. S1C**). Each effector handle design is associated with a DNA barcode on the same construct that will later be expressed as a circular RNA barcode^2^. By using degenerate DNA barcodes and subassembly procedures developed for massively parallel reporter assays^25,26^, we associate each effector handle design with multiple barcodes (**Fig. S2A**). Importantly, the resulting plasmid library is promoterless, such that its components cannot be expressed until after successful integration to a genomic landing pad via Bxb1-mediated recombination (**Fig. S1D**), as described further below.

**Figure 1.**
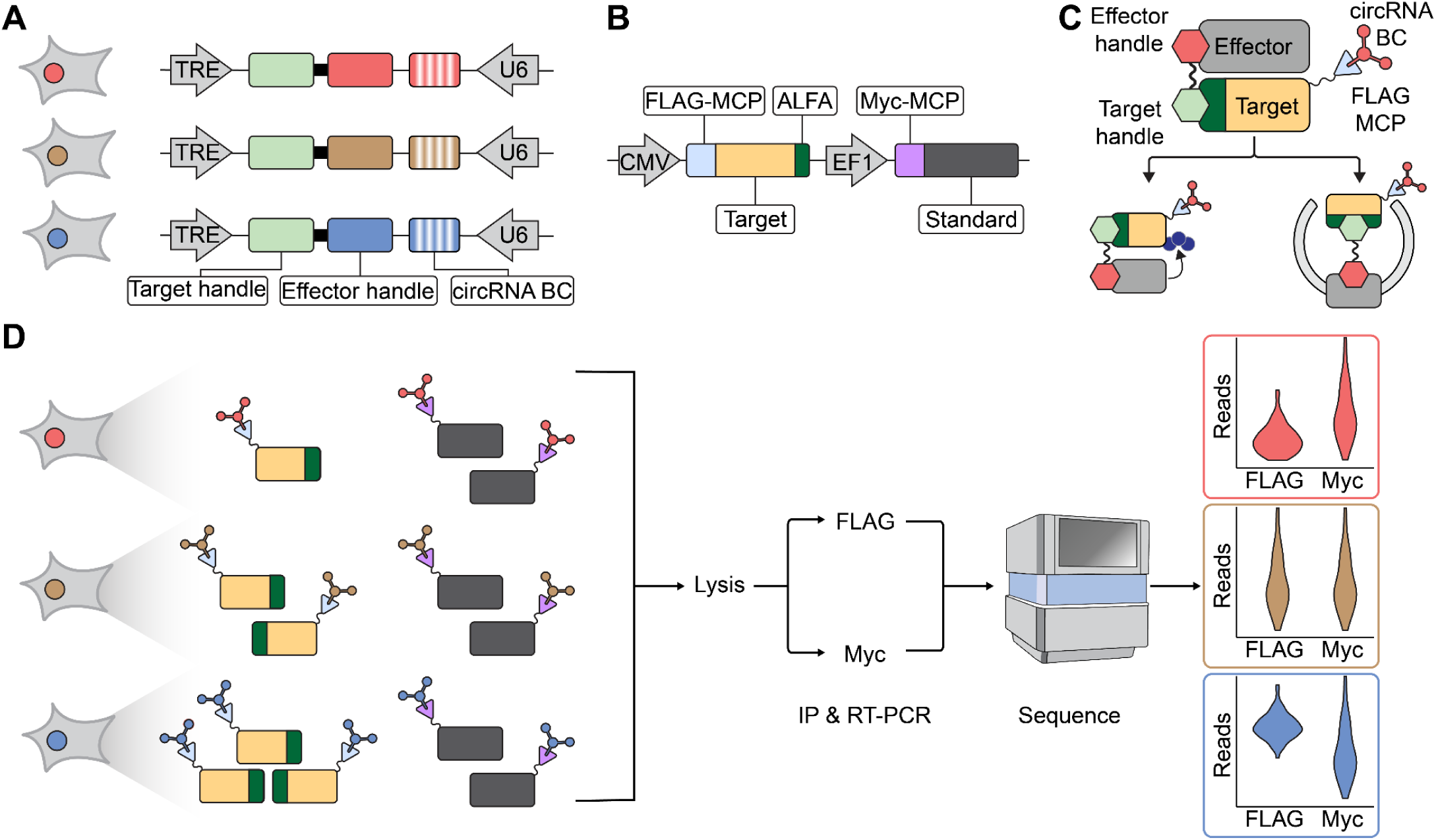
Multiplexed screening platform for *de novo* designed proteome editors using LABEL-seq. **A)** Schematic of genomic landing pad in engineered HEK293 cells following successful recombination. The locus encodes a doxycycline-inducible bifunctional proteome editor consisting of a constant target handle (green, NbALFA) fused to a variable effector handle generated through computational design (red, brown or blue; solid). For LABEL-seq^2^, the locus also encodes a constitutively expressed RNA species containing a barcode subsequence, MS2 hairpin, and terminal ribozymes (red, brown or blue; vertical stripes). Prior to recombination into the landing pad, each barcode subsequence is linked to a specific effector handle design via subassembly. Terminal ribozymes mediate circularization, generating a circular RNA barcode capable of binding the MCP domain via the MS2 hairpin. **B)** Schematic of the transiently transfected dual reporter construct. The construct consists of a CMV-driven target reporter (tan), which bears a FLAG-tagged MCP domain (pale blue) and ALFA epitope (dark green), and a EF1-driven standard reporter (dark grey), which bears a Myc-tagged MCP domain (purple). **C)** Schematic of a bifunctional proteome editor that successfully induces proximity of target and effector proteins. Also shown on the target protein is a FLAG-MCP domain (pale blue triangle) bound to a circRNA barcode that identifies the effector handle. Proximity induction can result in target degradation via ubiquitination (left, navy blue circles) or autophagy (right). **D)** Schematic of experimental workflow. Recombined HEK293 cells, each expressing one bifunctional proteome editor and one circular RNA barcode, are transiently transfected with the dual reporter construct. Design-identifying barcodes label both the target and standard reporters. Interaction of the editor with the target protein may result in its degradation (red), no effect (brown) or stabilization (blue). However, editors are not expected to interact with the standard protein. After 48 hours, cells are lysed, and the resulting lysate split to anti-FLAG and anti-Myc IPs. Co-immunoprecipitated circular RNA barcodes are subjected to RT-PCR and massively parallel sequencing. Effector handle designs that successfully promote degradation (red) or stabilization (blue) of the target protein are expected to shift the distribution of barcode counts in the anti-FLAG IP (target-specific) relative to the baseline defined by the anti-Myc IP (standard-specific), while those that have no effect are not (brown).

A second key component is a dual reporter construct encoding the target protein and a “standard” protein (**Fig. 1B**). The target protein is fused to a FLAG epitope (for IP), an MCP RBD (to recruit the LABEL-seq barcode) and an ALFA epitope (to recruit the bifunctional protein editor). The standard protein is tagged with a Myc epitope and the MCP RBD, but lacks any target recruitment epitope. In the LABEL-seq framework^2^, the Myc-tagged standard serves as an internal control. In any given cell, both the target and standard protein are bound by the same RNA barcode via the MS2-MCP interaction. Thus, target-associated barcode counts (FLAG pull-down) can be normalized to standard-associated barcode counts (Myc pull-down), isolating design-dependent changes in target abundance from technical or cell-to-cell variation.

We used Bxb1-mediated recombination to introduce the library of barcoded proteome editors into engineered HEK293 cells bearing a single copy of a landing pad^2^. Prior to recombination, the landing pad consists of an attP site sandwiched between convergent U6 and tetracycline response element (TRE) promoters (**Fig. S1D**). Successful recombination positions the proteome editor component downstream of the doxycycline-inducible TRE promoter, and the RNA barcode component downstream of the constitutive U6 promoter (**Fig. S1E**). Following selection for successful landing pad integrants, expression of the proteome editor is induced with doxycycline, and the dual reporter concurrently introduced via transient transfection.

At this stage, each successfully transfected cell should contain: (**i**) a single bifunctional proteome editor with a fixed target handle (NbALFA) and a variable (designed) effector handle; (**ii**) a single circular RNA species, bearing an MS2 hairpin and a barcode associated with the *cis-*encoded design; (**iii**) Target protein, bearing FLAG epitope, MCP domain, and ALFA epitope; and (**iv**) Standard protein, bearing Myc epitope and MCP domain. Within each cell, the MS2-bearing circRNA binds the MCP domain of both target and standard proteins, effectively barcoding individual protein molecules^2^. The levels of circRNA-barcoded target proteins may be modulated by successful effector handles, *e.g.* by recruiting them to proteasomes or lysosomes (**Fig. 1C**), while those of standard proteins should not. Forty-eight hours after editor induction and reporter transfection, the cells are lysed and the lysate split to either an anti-FLAG or anti-Myc IP (**Fig. 1D**). As per the LABEL-seq protocol, circRNAs recovered from each IP are then subjected to RT-PCR and massively parallel sequencing.

From the resulting sequencing data, editor-associated barcodes are counted, and their proportional contributions to the anti-FLAG IP-derived library and the anti-Myc IP-derived library are calculated and ranked. In this *trans* configuration setup of LABEL-seq, the bifunctional proteome editors whose associated barcodes are relatively enriched in the anti-FLAG IP-derived library can be inferred to have promoted stabilization of the target protein, while those whose associated barcodes are relatively depleted can be inferred to have promoted its degradation (**Fig. 1D**).

### *De novo* protein design of candidate effector handles

While developing this experimental framework, we also sought to select effector proteins for a proof-of-concept, and to computationally design binders to them as our candidate effector handles for bifunctional proteome editors. A recent proteome-wide study systematically profiled endogenous human proteins for their capacity to mediate proximity-dependent protein degradation or stabilization^19^. Guided by these results, we selected seven representative effectors associated with the major protein degradation pathways (**Fig. 2A**). Specifically, we chose FBXL12 and TRAF2 as components of distinct E3 ubiquitin ligase complexes, UCHL1 and USP38 as deubiquitinases (DUBs), and GABARAP, GABARAPL2, and MAP1LC3A as autophagy-related proteins (ATGs). In the cited study^19^, FBXL12 and the three ATGs were classified as degraders, while TRAF2 and the two DUBs were classified as stabilizers.

**Figure 2.**
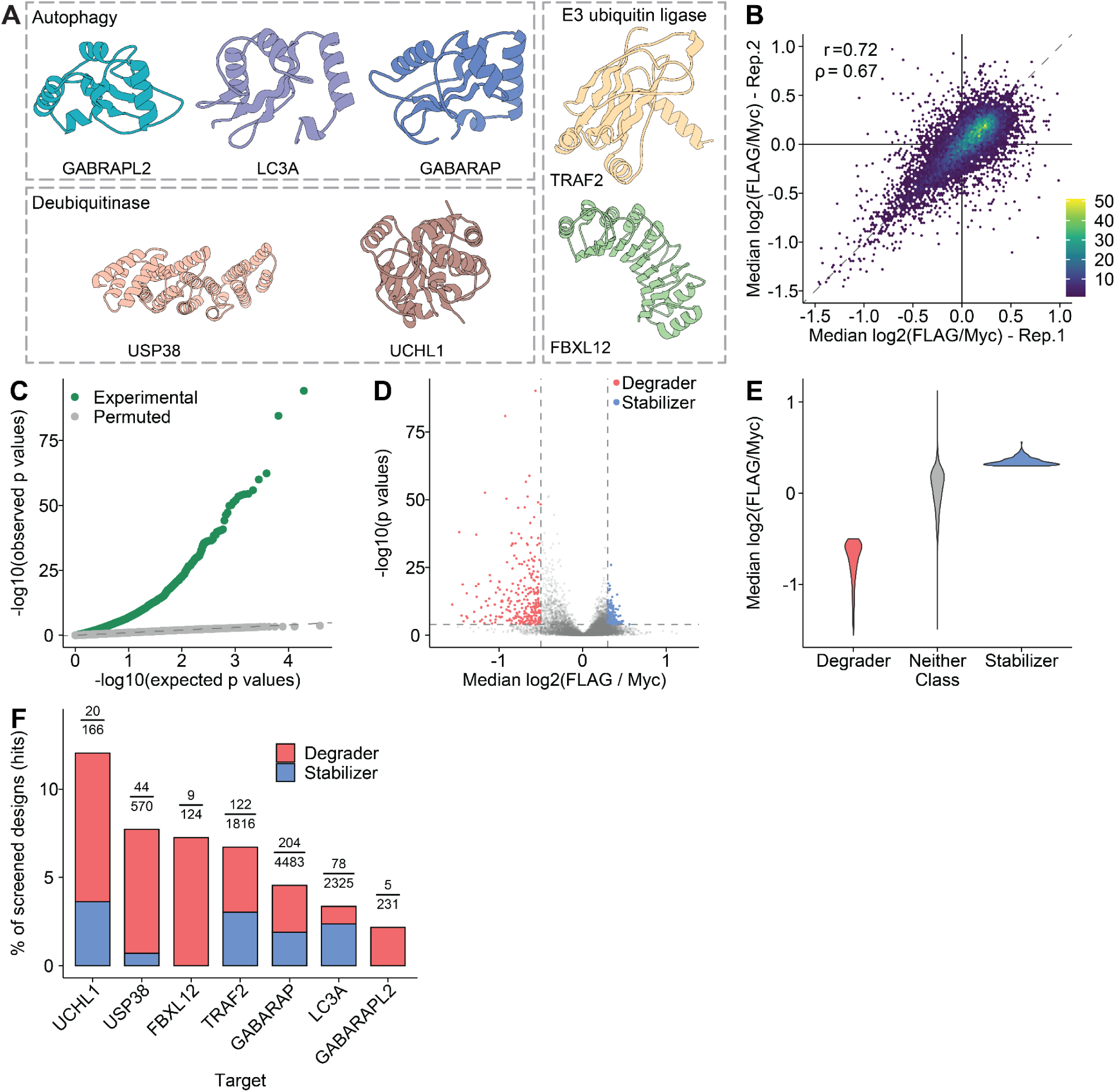
Identification of functional effector handle designs with LABEL-seq. **A)** Structures of effector proteins against which *de novo* protein binders were computationally designed. **B)** Hexagonal bin plot comparing the median log2(FLAG/Myc) fold-change for each design between two replicates. **C)** Quantile-quantile plot showing excess of low p-values (green points) in observed data, but no such excess when binder-barcode associations are permuted (grey points). Dashed line corresponds to null (x=y). **D)** Volcano plot showing effect sizes (*x*-axis; median log2(FLAG/Myc) fold-change) and significance (*y*-axis; −log10(p-value)) for each effector handle design. **E)** Violin plot showing effect distributions for effector handle designs classified as degraders (red; n = 277; *p* < 0.05 & median log2(fold-change) < −0.5), stabilizers (blue; n = 204; *p* < 0.05 & median log2(fold-change) > 0.3), or neither (gray). **F)** Barplot showing the percentage of binders classified as hits (y-axis) for each of 7 effector targets. Each bar indicates the percentage of hits classified as degraders (red) or stabilizers (blue). Above each bar, we also list the fraction on which the percentage was calculated (number of hits over number of designs for that effector).

To design candidate binders for these seven effectors, we employed deep-learning-based *de novo* protein design. First, RF Diffusion^15^ was used to build protein backbones predicted to engage each effector. We then used Protein Message Passing Neural Network (Protein-MPNN)^16^ to propose amino acid sequences for each backbone. Finally, the resulting candidates were structurally evaluated with AlphaFold2 (AF2)^27^. For each effector, we performed two iterative rounds of partial diffusion and sequence resampling to optimize binder–effector interfaces, and moved forward only with designs that passed AF2 confidence thresholds for interface predicted aligned error (PAE) and predicted local distance difference test (pLDDT) scores.

Altogether, we designed 10,225 candidate effector handles across seven effector targets, ranging from 125 for FBXL12 to 4632 for GABARAP (**Fig. S2B**; **Tables S1**, **S6**). Each design was 100 amino acids in length. These sequences were synthesized as a pool of oligonucleotides (Twist Biosciences), PCR-amplified, and cloned into a plasmid backbone. A 16-nucleotide (nt) degenerate barcode was then introduced downstream of each construct, generating the barcoded library that was ultimately recombined into the genomic landing pad (**Fig. 1A**). A high-confidence dictionary of mappings of barcodes to designs was generated by subassembly^25^ (**Figs. S2A**,**C-D**). Because the library was generated and maintained at high complexity, each effector handle was represented by multiple unique barcodes (mean 43 ± 41; **Fig. S2E**; **Table S2**). This is a key feature, as each barcode mitigates against technical variation by providing a quasi-independent internal replicate for a given design within the context of a single multiple experiment.

### ∼10,000-plex functional assessment of bifunctional proteome editors

This barcoded library—encoding 10,225 bifunctional proteome editors, each composed of a constant target handle and a variable, computationally designed effector handle—was introduced into HEK293 landing pad cells via Bxb1-mediated recombination (**Fig. 1A**; **Fig. S1C-E**). Following selection for successful recombinants, cells were split and transfected in duplicate with the dual reporter construct (**Fig. 1B**), and simultaneously treated with doxycycline (dox) to express the proteome editors. We chose glucokinase (GCK) as the target protein for this proof-of-principle experiment, primarily because GCK is monomeric, moderately sized (52 kDa) and cytoplasmically localized.

Replicate cultures were harvested, lysed, and divided for two immunoprecipitations (IPs): anti-FLAG to enrich the target protein and anti-Myc to enrich the standard protein (**Fig. 1D**). Circular RNA barcodes co-immunoprecipitated in each IP were reverse-transcribed, PCR-amplified, and subjected to massively parallel sequencing (**Fig. 1D**). To quantify the effects of individual editors on target protein abundance, barcode counts were first tabulated for each of the four samples (two biological replicates × two IPs; **Table S3**). For each barcode–design pair in the barcode dictionary, read counts were normalized to total reads per sample to obtain a per-sample read proportion for each barcode.

A key technical concern is whether recombination to the landing pad bottlenecked the complexity of the original library. To test this, we tallied the number of barcodes associated with each design. Reassuringly, “summary read proportions” were highly correlated between the plasmid library and the experimental libraries (Pearson’s *r* = 0.98, Spearman’s ⍴ = 0.98). However, ∼2% of designs exhibited substantial “dropout” when introduced to cells (**Fig. S3A**). This phenomenon was consistent between replicates, but differed across effector targets (**Fig. S3B**). For example, effector handles designed to bind DUBs were 8.5-fold more likely to cause >50% dropout than those designed to bind other effector targets (10.2% of DUB binders [79/776] vs. 1.2% [110/9,200] of other binders). We speculate that these designs are toxic to HEK293 cells when highly expressed, either intrinsically or through their interaction with effectors.

After excluding barcodes with low representation (<10⁻⁷ of reads in any of the four samples) and then designs represented by fewer than three barcodes, 9,715 designs (95%) were retained for further analysis. The effects of effector handle designs on the ratio of FLAG-to-Myc read proportions spanned a three-fold range and were reproducible between technical replicates (**Fig. 2B**; comparing the median ratio across all barcodes associated with a given design in Rep-1 vs. Rep-2: Pearson’s *r* = 0.72, Spearman’s ⍴ = 0.67; **Table S4**). Across the two replicates, there were 3,699 effector designs that significantly stabilized or destabilized the target protein (*p* < 0.05; Wilcoxon signed-rank test in which the FLAG vs. Myc read proportion rank distributions are compared for barcodes associated with a given design; *p* values were corrected for multiple-hypothesis testing using the Benjamini-Hochberg procedure). Among these, 2,016 (55%) were significant in both replicates, with perfectly concordant directional effects (2,016/2,016 [100%]). This directional concordance overwhelmingly extended to designs significant in only one replicate (1,575/1685 [94%]). Encouraged by these results, we merged the sets of design-associated Flag and Myc read proportions observed in the two replicates (**Table S5**). In the combined data, 55% (5,299/9,715) of designed effector handles significantly stabilized or destabilized the target protein (*p* < 0.05; Wilcoxon signed-rank test; *p* values corrected using the Benjamini-Hochberg procedure).

Although we are using conservative procedures in testing for statistical significance (non-parametric test; hypothesis control), we performed an additional check in which we simply permuted the barcode-design assignments and recalculated significance by the same procedures. In the resulting quantile-quantile plot, we observe a massive excess of small *p-*values with the true barcode-binder assignments, while the permuted barcode-binder assignments result in a distribution of *p-*values that closely tracks expectation (**Fig. 2C**).

Overall, these analyses show that most designed effector handles (**Table S6**) exhibit significant, reproducible, directionally consistent effects on the levels of the target protein. However, most of these effects are small in magnitude. For our further investigations, we focused our attention on the 481 bifunctional proteome editors with the largest effect sizes, including 277 “degraders” (*p* < 0.05 & median log2(fold-change) < −0.5) and 204 “stabilizers” (*p* < 0.05 & median log2(fold-change) > 0.3) (**Fig. 2D**). With these thresholds, functional designs were obtained for both the UPS (130 degraders, 64 stabilizers) and autophagy (147 degraders, 140 stabilizers) pathways. Of note, degraders were more likely to have stronger effect sizes than stabilizers, *e.g.* 35 degraders, but no stabilizers, altered target protein abundance by >2-fold (**Fig. 2E**).

Our “hit rate” in terms of whether designs would succeed as either degraders or stabilizers varied substantially by effector, from 2% for GABARAPL2 (5/231) to 12% for UCHL1 (20/166) (**Fig. 2F**). As noted above, FBXL12 and the three ATGs were previously classified as degraders, while TRAF2 and the two DUBs were classified as stabilizers^19^. However, only FBXL12 and GABARAPL2 were fully concordant with this prior. For example, 9/9 successful FBXL12 binders were degraders rather than stabilizers. In contrast, successful designs for the other effectors included both degraders and stabilizers (**Fig. 2F**). For example, we expected binders that recruited the target protein to either of the two DUBs (UCHL1 and USP38) to increase its levels. However, not only did these two effectors have the highest “hit rates” (12% and 8%, respectively), their associated designs were mostly degraders (14/20 [70%] and 40/44 [91%], respectively).

### Singleton validation of bifunctional protein degraders and stabilizers

We next sought to validate selected bifunctional protein degraders and stabilizers identified in the ∼10,000-plex cellular screen. We chose not to pursue further validation of designed effector handles targeting GABARAPL2 due to the paucity of designs exhibiting large effect sizes. For the remaining six effectors, we selected the two degraders and one stabilizer for singleton validation experiments (**Fig. S4**; **Table S7**). As a negative control, we also selected a binder that consistently exhibited no effect on the levels of the target protein. We then generated constructs encoding a constitutively expressed NbALFA (target handle; C-terminus) fused to one of these 19 binders (effector handle; N-terminus). These constructs also express mTagBFP2 from an independent EF1 promoter as a transfection marker.

To measure protein abundance changes in an assay orthogonal to the original LABEL-seq screen, we employed a bicistronic GFP-IRES-mCherry protein abundance reporter that enables ratiometric quantification of GFP-fusion protein abundance normalized to mCherry. We generated two dox-inducible variants of this reporter. The first reporter encoded a FLAG-GFP-GCK-ALFA-IRES-mCherry-2A-Puro cassette. As a negative control, we sought to abolish the interaction between the binder and the ALFA-tagged GFP protein. To accomplish this, a second version of the reporter was cloned in which the ALFA tag was deleted in-frame, leaving FLAG-GFP-GCK only. Stable polyclonal HEK293T lines expressing either reporter were generated via piggyBac transposition of the reporter plasmids. Effector handle-encoding plasmids were transiently transfected into each reporter line in triplicate. Reporter expression was induced with dox for 24 hrs post-transfection, and then the GFP:mCherry ratios of mTagBFP2-positive cells were assessed by flow cytometry at 48 hrs post-induction.

Within each reporter line, GFP:mCherry ratios measured in the presence of each protein editor (*i.e.* NbALFA fused to an effector handle) were normalized to the mean GFP:mCherry ratio of the negative control binder (**Fig. 3A-B**, grey points). For 12/12 degraders, we observed ratios consistently below that of the no-effect control (**Fig. 3A**, red points), while for 5/6 of the stabilizers, we observed ratios consistently above that of the no-effect control (**Fig. 3A**, blue points). Altogether, 17/18 (94%) effector handle designs subjected to singleton validation exhibited directional concordance with expectation. The absence of the ALFA tag in the GFP:mCherry reporter abolished the effects of these effector handles, providing further evidence of their specificity (**Fig. 3B**). These effects can also be visualized as a slope plot to more directly compare the GFP:mCherry ratios induced by each binder in each cell line (**Fig. 3C**). The GABABRAP-D1 effector handle induced the strongest degradation effect, with a mean normalized GFP:mCherry ratio of 0.26 indicating a 3.9-fold decrease in abundance of the reporter protein. TRAF2-S1 induced the strongest stabilization effect, a 1.4-increase in levels of the reporter protein.

**Figure 3.**
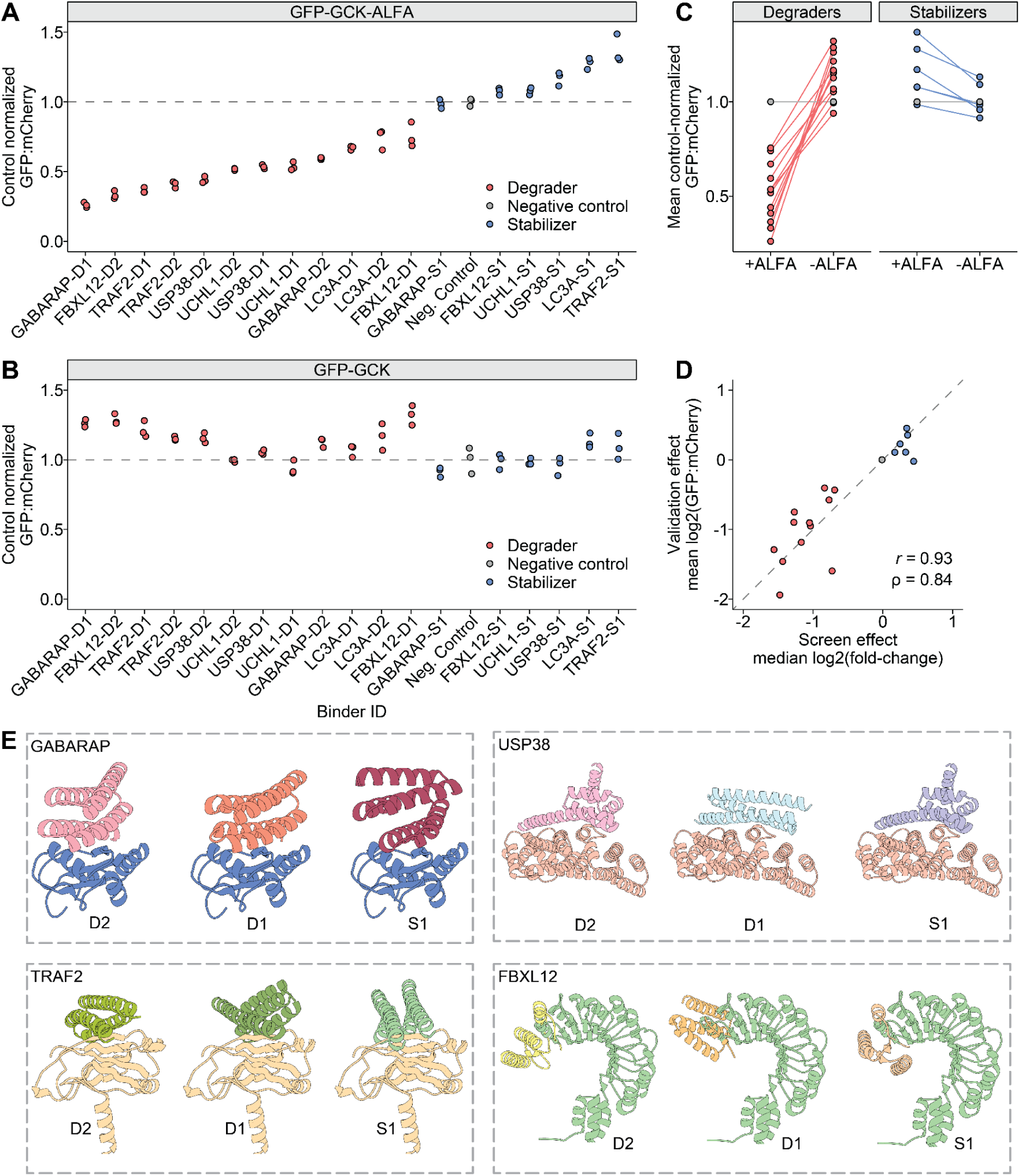
Singleton validation of functional designed effector handles. **A)** Dot plot of control-normalized GFP∶mCherry ratios measured in cells expressing a EGFP-GCK-ALFA reporter and indicated candidate effector handle designs. Each point represents a biological replicate and the dashed line at 1 indicates the mean of the negative-control effector handle (grey points). Effector handles are ordered along the *x*-axis by mean normalized GFP:mCherry ratio, Dx and Sx (where x is a numeral) indicate degraders and stabilizers, respectively. **B)** Same as in panel **A** but measured in cells expressing a EGFP-GCK reporter lacking the ALFA tag. Effector handles in panel **B** maintain the same left-to-right ordering as in panel **A**. **C)** Slope plot faceted by effector class showing mean GFP:mCherry ratios measured for each binder in each cell line. Binder measurements in each line are connected with a line to highlight the effect of deleting the ALFA tag from the reporter construct. **D)** Scatterplot comparing measured effects from the ∼10,000-plex screen and LABEL-seq (*x*-axis) vs. the singleton validations (*y*-axis). **E)** Visualizations of predicted structural models for indicated effector-binder pairs. Identifiers in each panel (*i.e.* D2, D1, S1) are matched to identifiers in panels **A** and **B**.

We also compared effects estimated from the ∼10,000-plex experiment with LABEL-seq against effects measured via the orthogonal GFP:mCherry reporter assay, and found these to be highly concordant (Pearson’s *r* = 0.93, Spearman’s ⍴ = 0.98; **Fig. 3D**). Finally, we extended these validations from HEK293T cells to HepG2 cells and once again observed strong concordance (Pearson’s *r* = 0.91, Spearman’s ⍴ = 0.93; **Fig. S5**), suggesting that these bifunctional proteome editors may function in all cell types where the effector proteins to which they bind are active.

Finally, we then examined the design models of binder-effector complexes for effector handles that induced >50% degradation (**Fig. 3E**). Designs against GABARAP, FBXL12, USP38, and TRAF2 each targeted a conserved binding interface within their respective targets. The highly performing degrader targeting FBXL12 overlaps with the C-terminus of FBXL12, which is normally responsible for engaging native substrates for degradation. This observation suggests that the conformational flexibility of the C-terminus of FBXL12 may permit binder engagement and substrate redirection, consistent with the intended design principle of redirecting FBXL12 to degrade the protein recruited by the target handle.

### Targeting an endogenous protein with designed effector handles

We next asked whether designed effector handles could mediate degradation of endogenous proteins by replacing the ALFA-targeting NbALFA handle with a binder specific to an intracellular target. We focused on the anti-apoptotic BCL-2 family protein MCL1, which has a short half-life due to UPS-mediated degradation, is overexpressed in many cancers, and has previously been targeted by high-affinity designed binders^5^.

The top degrader design for each of six effectors (**Table S7**) was cloned into a lentiviral backbone as a bivalent fusion with an anti-MCL1 binder. MDA-MB-231 breast cancer cells were transduced with individual MCL1-degrader fusions, and endogenous MCL1 protein levels were assessed by Western blot 72 hrs post-transduction (**Fig. S6**). MCL1 was expressed at low levels in untransduced cells and in cells expressing a GFP-specific nanobody (NbGFP) (**Fig. S6**, lanes 1-2). Notably, MCL1 levels were elevated in cells expressing a negative control construct in which the MCL1 binder was fused to a non-functional effector handle from the LABEL-seq screen. This upregulation is consistent with prior studies showing that MCL1 inhibitors or binders can stabilize MCL1 by blocking native ubiquitination, leading to accumulation of non-functional MCL1 that is unable to sequester pro-apoptoptic factors^28–30^.

Accordingly, MCL1 levels were normalized to those observed with the negative control MCL1-non-functional binder fusion. Among the tested constructs, GABARAP and LC3A effector handles showed the strongest effects, reducing MCL1 protein levels to 50% and 27% of control levels, respectively (**Fig. S6**). These findings mirror the strong degradation effects of autophagy-related effector handles observed in our ∼10,000-plex experiment and singleton validation assays. In contrast, fusions incorporating UPS-related effectors showed only modest or stabilizing effects on endogenous MCL1, despite their activity in the GCK–ALFA reporter system, suggesting target- or cell line-specific dependencies. Testing several effector handles for each of these autophagy effector confirmed that the D1 designs for GABARAP and LC3A (*i.e.* the ones exhibiting the strongest effects in both the multiplex discovery and singleton validation experiments) also yielded the strongest activity against endogenous MCL1 (**Fig. 4A**).

**Figure 4.**
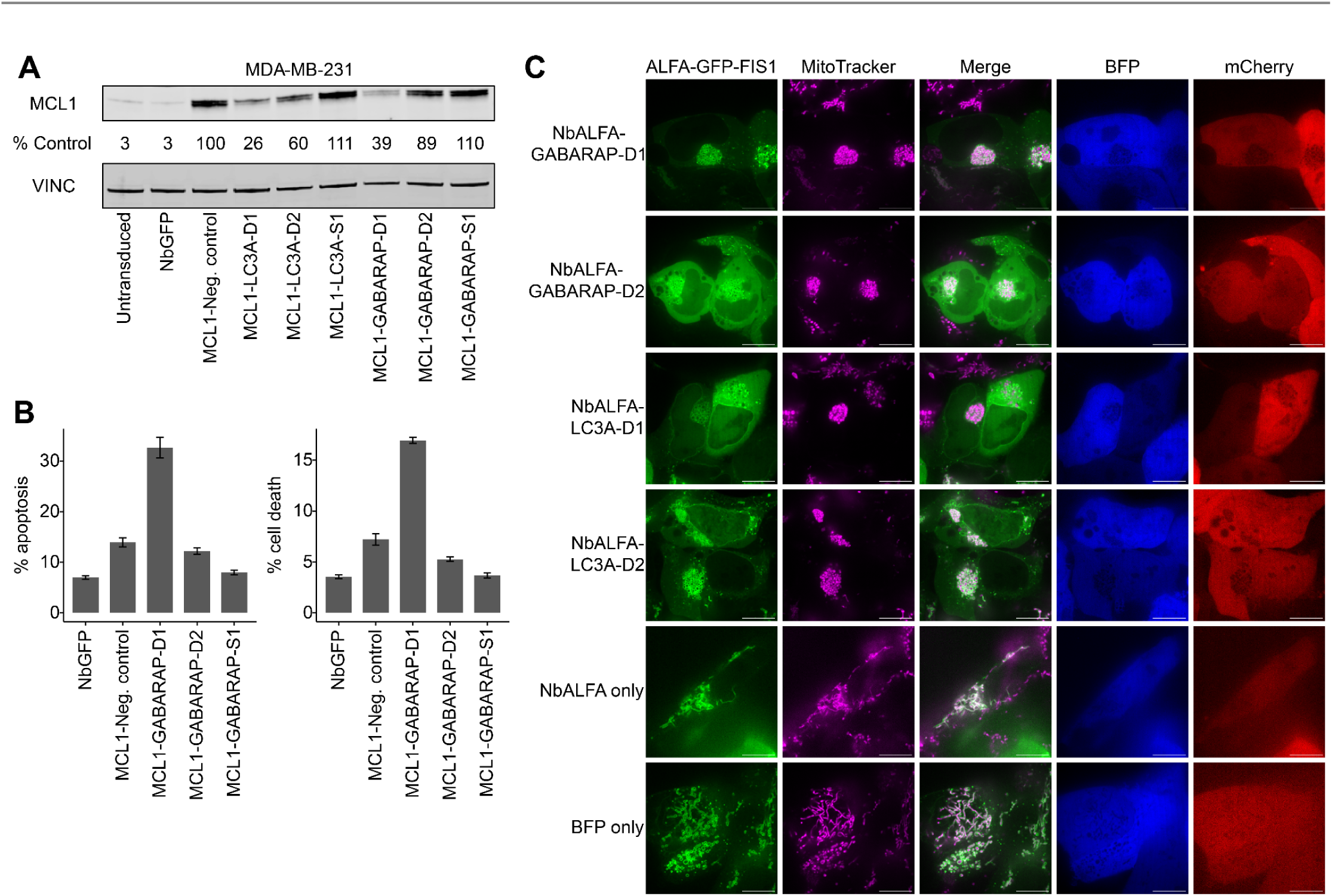
Designed autophagy effectors mediate endogenous protein degradation, promote cancer cell apoptosis, and perturb mitochondrial networks. **A)** Western blot analysis of MCL1 levels in MDA-MB-231 cells, 72 hours after transduction with indicated MCL1-effector binder fusions, as well as untransduced and GFP-nanobody transduced controls. **B)** Percentage of MDA-MB-231 cells positive for apoptotic (left) or cell death markers (right) as measured by flow cytometry, 72 hours after transduction with indicated MCL1-GABARAP binder fusions or controls. Error bars indicate ± SEM (n = 3). **C)** Confocal microscopy images of ALFA-GFP-FIS1 (green) and mitochondria (magenta). Images were taken 24 hours after co-transfection of HEK293T cells with plasmids encoding ALFA-GFP-FIS1 and indicated MCL1-effector binder fusions or controls. mCherry and BFP are transfection markers for the ALFA-GFP-FIS1 and MCL1-effector binder fusion plasmids respectively.

To evaluate the functional consequences of MCL1 degradation, we quantified apoptosis and cell death in MDA-MB-231 cells expressing GABARAP-targeting fusions. Consistent with their degradation effects, GABARAP-D1—which produced the strongest degradation by Western blot—also induced increased apoptosis and cell death, whereas non-degrading GABARAP-targeting fusions had no effect (**Fig. 4B**; **Fig. S7**). Previous studies have shown that the functional (anti-apoptotic) pool of MCL1, rather than total abundance, determines the apoptotic threshold^31^. The bifunctional proteome editor targeting MCL1 and GABARAP may therefore act through dual mechanisms: promoting degradation while simultaneously preventing sequestration of pro-apoptotic factors, thereby neutralizing MCL1’s protective function even when residual protein remains. These results indicate that degradation of the anti-apoptotic pool of MCL1 triggers apoptosis and cell death, consistent with prior findings from PROTAC-based degraders^30,32^. Collectively, these data demonstrate that de novo designed effector handles can drive degradation of an endogenous oncoprotein and elicit functional phenotypes in human cancer cells.

### Perturbation of mitochondrial organization

Beyond modulating individual proteins, we wondered whether designed binders could be used to redirect or reorganize subcellular structures. To test this, we performed live-cell imaging of HEK293T cells co-transfected with plasmids encoding ALFA–GFP–FIS1 and NbALFA–effector handle fusions, and visualized mitochondrial networks by confocal microscopy. FIS1 (mitochondrial fission 1 protein) localizes to the outer mitochondrial membrane; thus, ALFA–GFP–FIS1 allows visualization of mitochondria via the GFP signal (**Fig. 4C**, GFP channel, green). GFP fluorescence co-localized with mitochondria stained with MitoTracker (**Fig. 4C**, MitoTracker, magenta), confirming correct targeting.

Strikingly, cells co-expressing NbALFA fusions with GABARAP or LC3A effector handles displayed aggregated mitochondrial networks (**Fig. 4C**, top rows), in sharp contrast to the dispersed, reticular morphology seen in control cells co-expressing NbALFA alone or in BFP-only controls (**Fig. 4C**, bottom rows). Given the established roles of GABARAP and LC3A in autophagy, these aggregates likely reflect synthetic tethering or sequestration of mitochondria into autophagosomal compartments^7,8^. Although the precise mechanism remains to be determined, the binder-dependent clustering of mitochondria demonstrates that designed effector handles can act beyond single-protein modulation to reprogram organelle-scale organization, illustrating the potential for programmable subcellular remodeling.

## DISCUSSION

Here we describe a multiplex framework for quantifying the intracellular activities of *de novo* designed proteome editors. In this proof-of-concept, we evaluated ∼10,000 designed binders of components of the UPS and autophagy pathways as effector handles within bifunctional proteome editors. Remarkably, the majority of designed binders reproducibly altered the abundance of a target protein, although most effects were modest in magnitude. We validated a subset of the most active “degraders” and “stabilizers” in an orthogonal protein abundance assay, and found these exhibited strong concordance with the measurements obtained in the multiplex experiment, both in terms of directionality and magnitude of effects.

To date, the field of targeted protein degradation has been dominated by small molecules such as PROTACs. However, their development is constrained by the limited availability of well-characterized effector ligands and the requirement for chemical tractability of target proteins. By integrating *de novo* protein design with a multiplex functional screen based on LABEL-seq, we establish a general framework that circumvents these constraints. Computational design enables generation of bespoke binders to virtually any effector, target, or interface, while the multiplex assay provides a scalable means of measuring and optimizing their activities within living cells.

For this initial demonstration, we employed an ALFA-tagged GCK reporter to evaluate designed effector handles. Even for binders targeting the same effector interface, we observed a broad spectrum of effects on reporter abundance, suggesting that attributes such as binding affinity, dissociation rate, intracellular stability, or expression may modulate efficacy. Further studies will be necessary to elucidate the precise mechanisms and effector dependencies of our designed binders, both to validate their mode of action and to guide future iterations of design. Although we designed our study with the expectation that recruiting certain effectors would result in either degradation or stabilization, our results suggest a continuum of effects rather than a binary outcome. Furthermore, as our experiment produced quantitative measurements for hundreds to thousands of designs per effector, investigations of the biophysical features that determine the directionality and magnitude of these outcomes may be enabled by the data reported here.

GCK’s long half-life likely limited our dynamic range for detecting turnover, and more generally, the portability of designed effector handles across different targets remains an open question. Consistent with this, effector handles that recruited autophagy effectors (*e.g.* LC3A, GABARAP) successfully degraded the endogenous oncoprotein MCL1 when paired with an anti-MCL1 binder, whereas those targeting the UPS did not. These findings suggest that the efficacy of bifunctional proteome editors will depend on both the effector and the context of the recruited target.

Looking forward, this framework opens several avenues for scaling and diversification:

First, *expanding effector diversity* (**Fig. 5A**). The present study targeted only seven effectors, but the same approach could be extended to hundreds of components across degradation, stabilization, and trafficking pathways. Such expansion would yield a rich toolkit of effector handles for proteome editing across cell types and physiological contexts.

**Figure 5.**
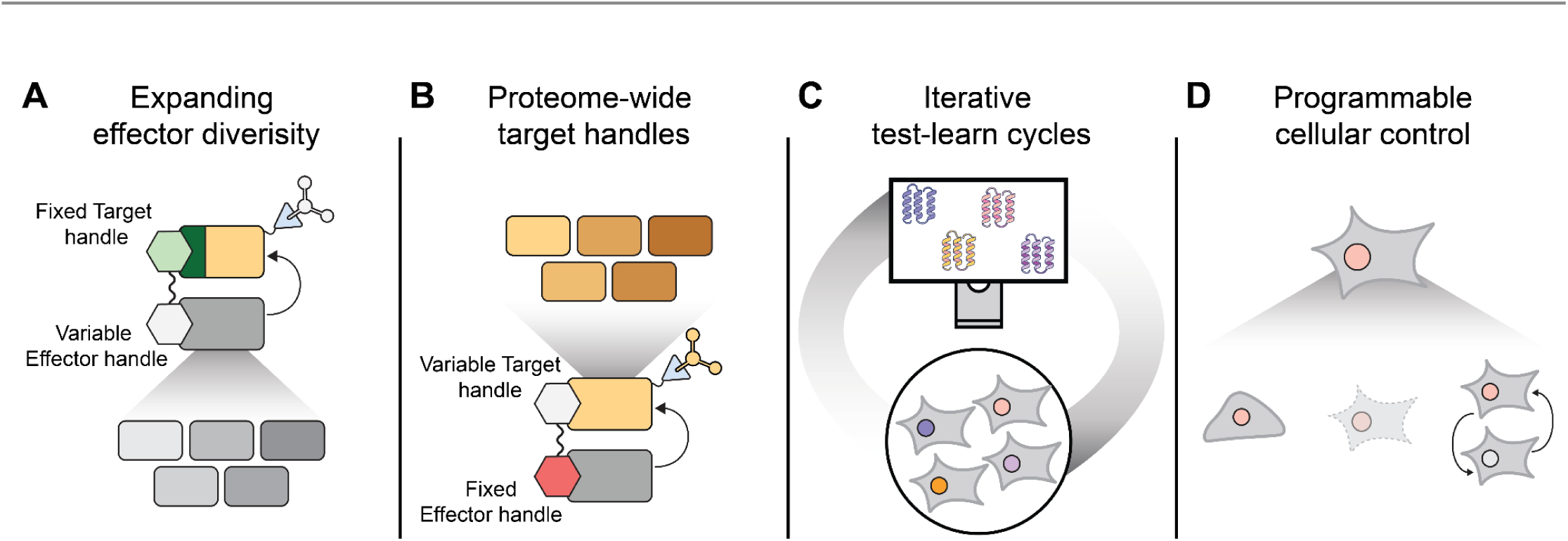
Potential applications of multiplex cellular screens of *de novo* designed proteins. **A)** A fixed target handle (green) can be fused to a library of designed effector handles with the goal of enabling diverse effector outcomes to be programmed. **B)** A fixed effector handle (red) can be fused to a library of designed target handles with the goal of broadening the set of recruitable proteins to the entire human proteome. **C)** Iterative cycles of effector or target handle design and experimental characterization may yield data which serves the improvement of the design of intracellularly functional binders. **D)** Installation of conditional control of protein editors may enable programmable cellular control. Potential applications include programming inducible changes to target protein abundance or localization, cell state, cell death, and cell-to-cell communication.

Second, *systematic design and validation of target handles for the entire human proteome* (**Fig. 5B**). By fixing a validated effector handle and instead varying the target handle, large-scale intracellular screens could identify binders for virtually every human protein). In principle, this would create a comprehensive “phone book” of functional binder pairs—enabling the programmable recruitment of any protein to any effector, location, or function—analogous to how base-pairing rules empower genome editing.

Third, *mechanistic and modeling advances* (**Fig. 5C**). Unlike traditional *in vitro* binding assays, our assay measures designs in the native cellular milieu and yields quantitative functional data for thousands of designs in a single experiment. These datasets, coupled with structural models, provide fertile ground for understanding how designed interactions translate to function within cells and for improving next-generation generative models for binder design.

Fourth, *extending proteome editing beyond abundance* (**Fig. 5D**). Beyond modulating protein levels, bifunctional proteome editors could be adapted to control a broader range of protein and cellular phenotypes, *e.g.* as we have shown here by perturbing mitochondrial networks. The same multiplex framework could also incorporate conditional regulation—via small molecules, light, or engineered feedback—to achieve tunable and context-dependent control of intracellular processes. Together, these extensions would transform proteome editors from degraders or stabilizers into general-purpose actuators for programmably reshaping cellular function.

Collectively, these advances point toward a future in which protein design and multiplex functional screening converge into a programmable toolkit for remodeling the intracellular proteome and the cell itself.

---

**Figure S1.**
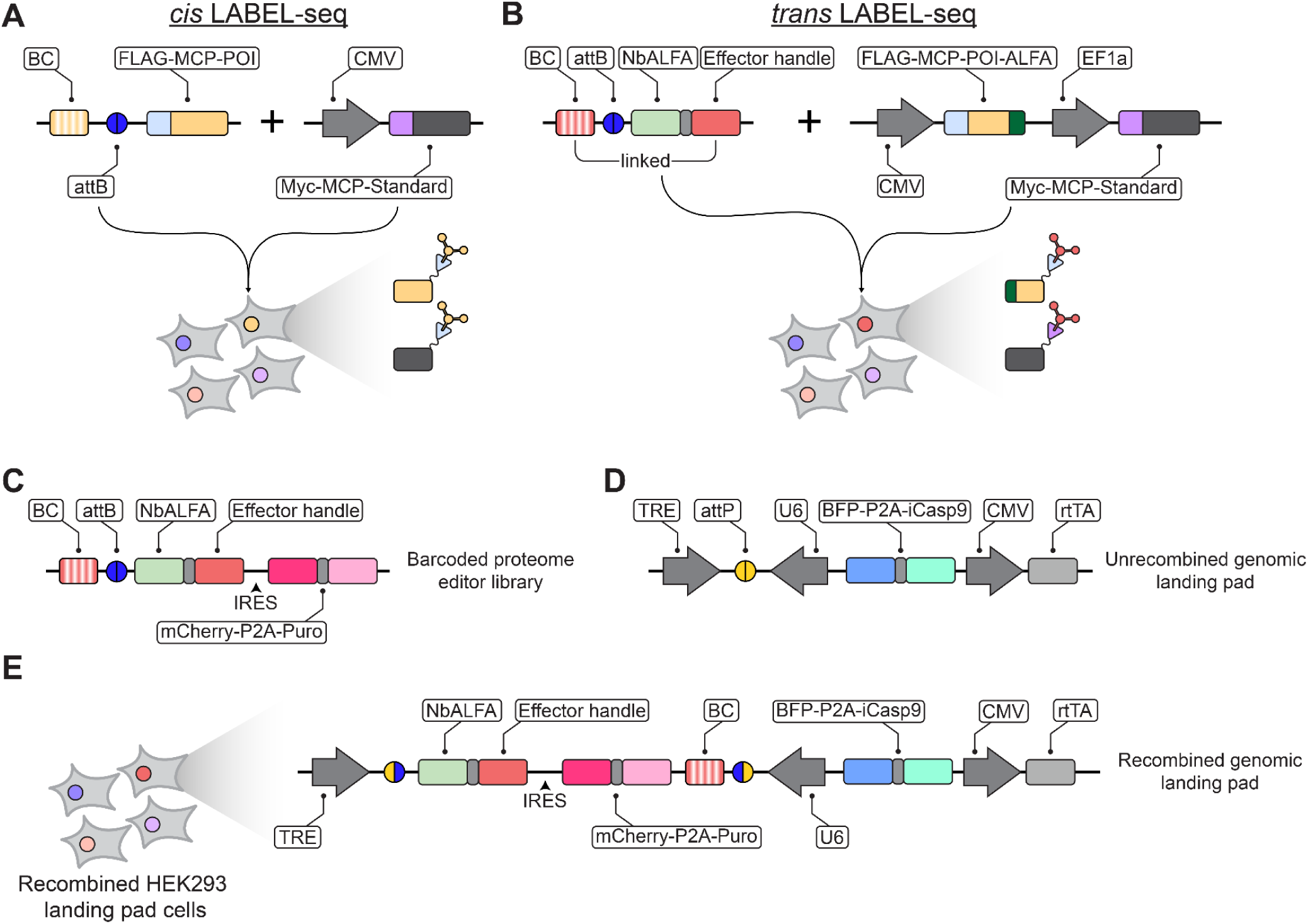
Schematic of genomic landing pad and *cis* versus *trans* LABEL-seq assays. **A)** Diagram of *cis* LABEL-seq in which the barcode (BC) and FLAG-MCP-POI protein are encoded from the same DNA molecules. **B)** Diagram of *trans* LABEL-seq in which barcode (BC) and FLAG-MCP-POI-ALFA protein are encoded from distinct DNA molecules. **C)** A barcoded proteome editor library prior to recombination at the landing pad. The barcoded proteome editor library also has a mCherry-P2A-Puro cassette that is translated from an upstream internal ribosome entry site (IRES). **D)** Diagram showing the single-copy genomic landing pad prior to recombination, as well as a barcoded element library. The landing pad contains an attP site flanked by convergent tetracycline response element (TRE) and U6 promoters, a BFP-P2A-iCasp9 cassette, and a CMV-driven reverse tetracycline transactivator (rtTA). A barcoded element library with an attB site can be recombined with the genomic attP site via Bxb1-mediated recombination. **E)** Diagram showing the single-copy genomic landing pad following recombination, which places the library element and IRES-mCherry-P2A-Puro cassette under control of the TRE promoter and the circular RNA barcode under control of the U6 promoter.

**Figure S2.**
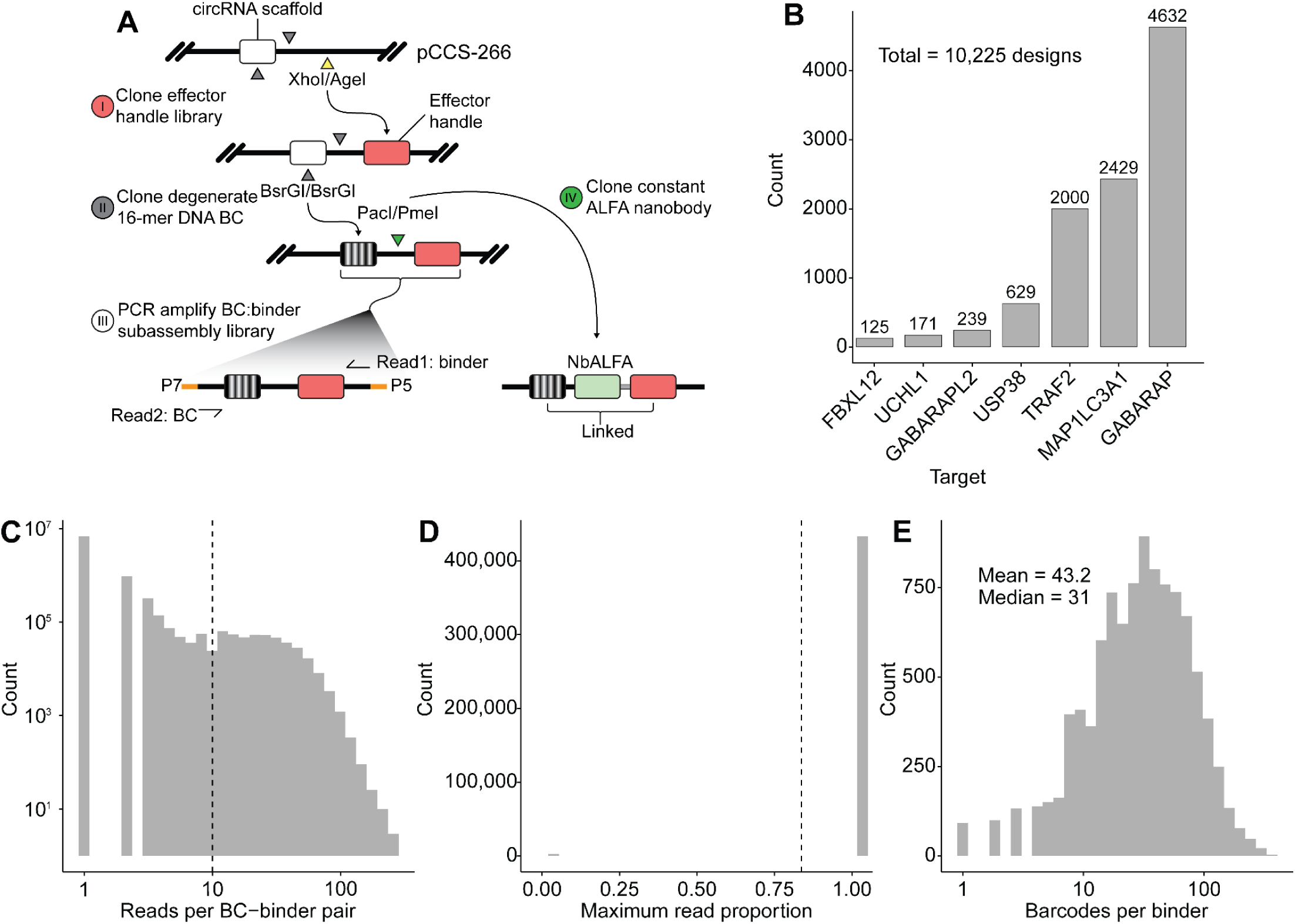
Plasmid library composition and characterization. **A)** Schematic detailing how the barcoded binder library was cloned, including subassembly. **B)** Barplot showing the number of designs generated for each target effector. **C)** Histogram showing the distribution of the number of sequencing reads per barcode-binder pair. The vertical dashed line indicates the threshold above which barcode-binder pairs were considered valid. **D)** Histogram showing the maximum read proportion of individual barcodes. For an individual barcode, the proportion of reads associated with any binder was computed to ascertain the fidelity of barcode binder association. The vast majority of barcodes were associated with a single binder. **E)** Histogram showing the distribution of the number of barcodes associated with each binder.

**Figure S3.**
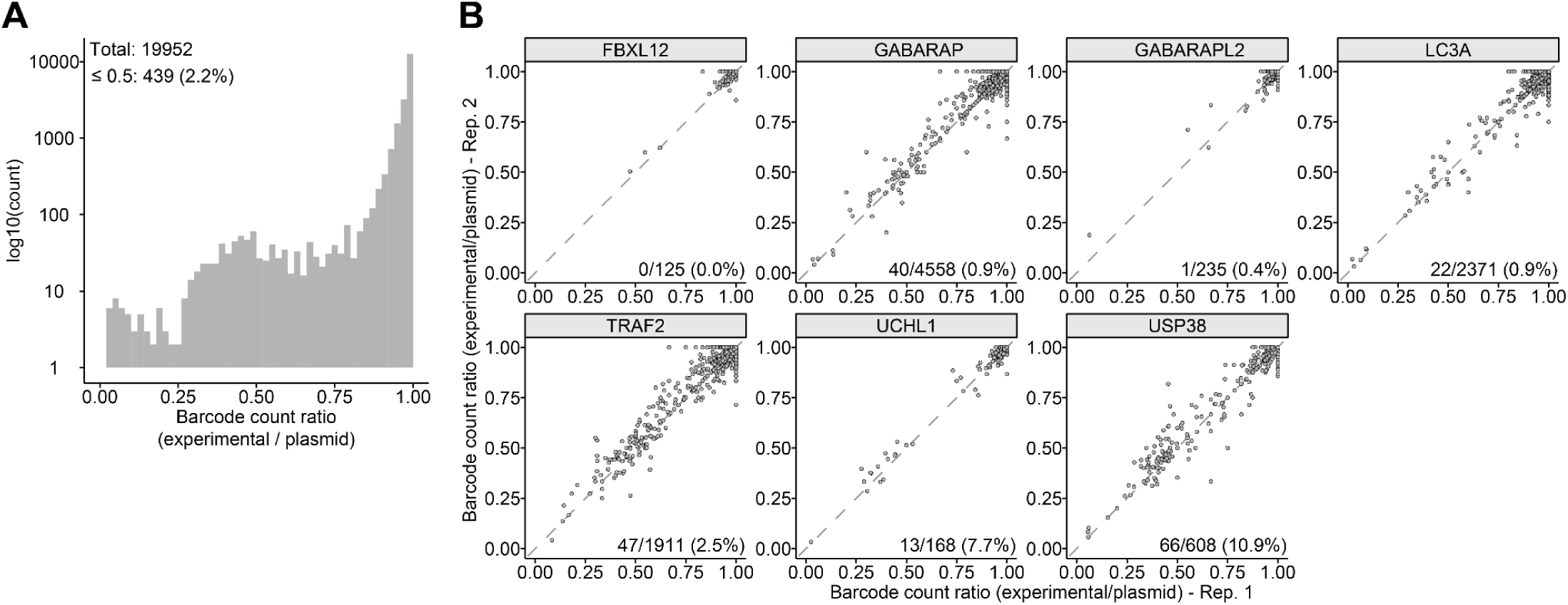
Comparison of plasmid and experimental barcodes. **A)** Histogram showing distribution of the ratio of experimentally recovered barcodes to plasmid library barcodes for each binder across two transfection replicates. The total number of binders represented in the plot is indicated, as well as the number and percentage of binders with a ratio ≤ 0.5 in either of the two replicates. The distribution suggests that some binders are “dropping out” of the experiment when expressed in cells. **B)** Scatter plots comparing the ratio of experimentally recovered barcodes to plasmid library barcodes for each binder between the two transfection replicates. Each point represents an individual binder. For each effector target, the number of binders with a ratio ≤ 0.5 in both replicates is indicated, as well as the percentage. The patterns suggest that the drop-out phenomenon is reproducible between replicates, and moreover is more likely to occur with binders designed against some effectors (*e.g.* USP38; 10.9%) than others (*e.g.* FBXL12; 0%).

**Figure S4.**
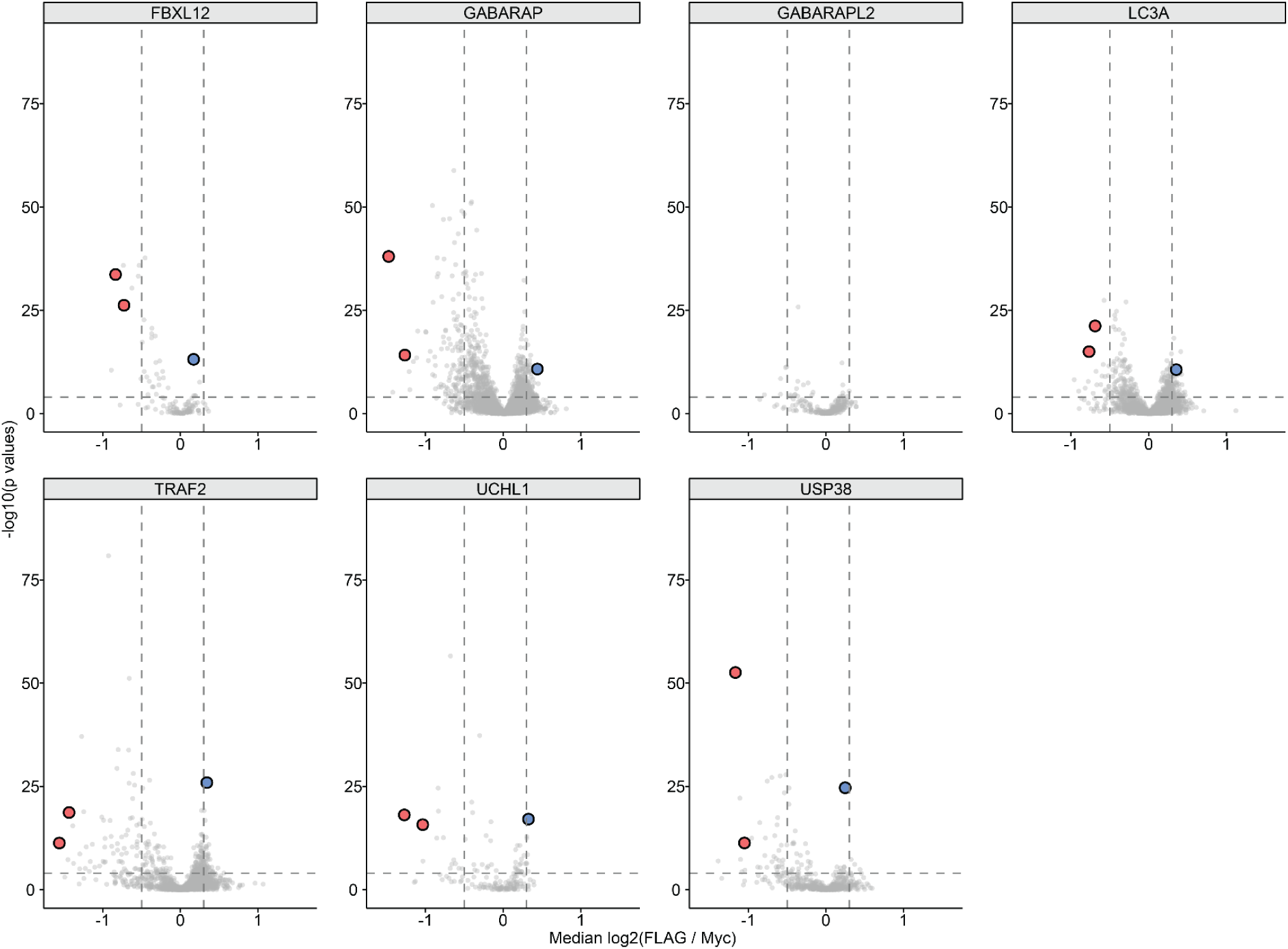
Assessment of hit rate per effector and selection of effector handles for validation experiments. Volcano plots comparing the distribution of effect sizes (x-axes) and significance (y-axes) faceted by effector. Degrader (red points) and stabilizer (blue points) effector handles selected for validation are indicated.

**Figure S5.**
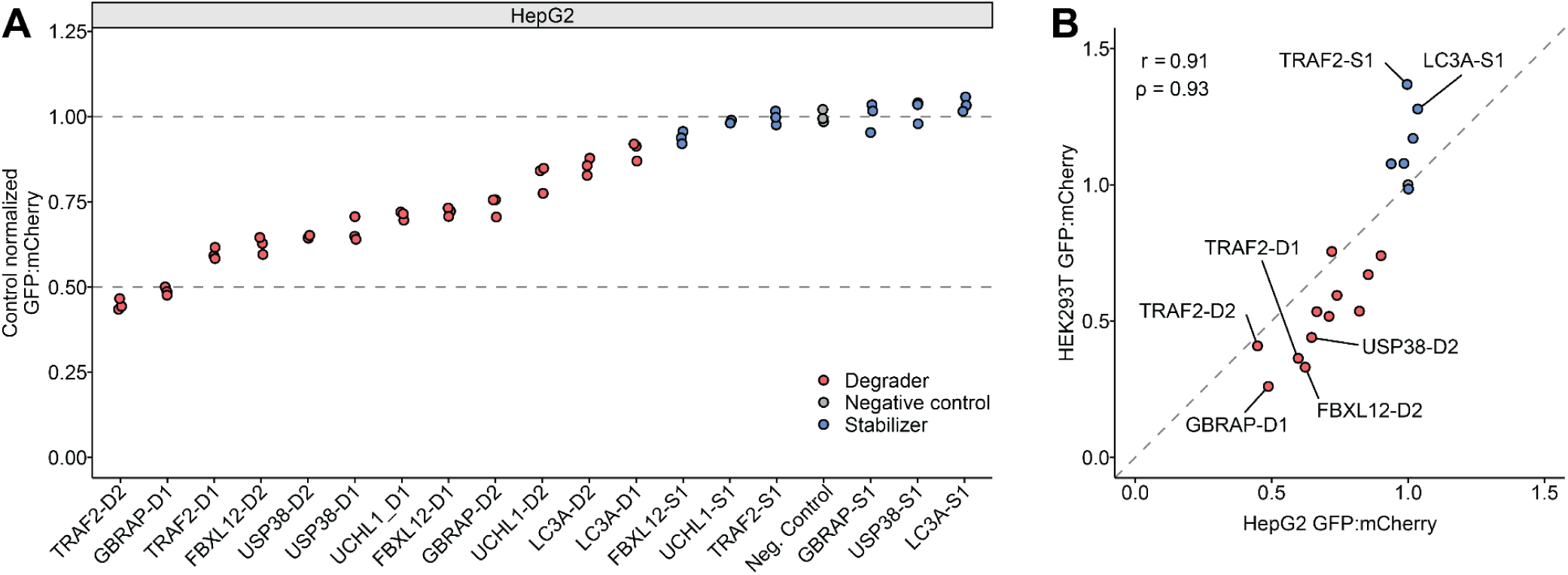
Binder-induced GCK degradation validates in an alternative cell line. **A)** Dot plot of control-normalized GFP∶mCherry ratios measured in HepG2 cells expressing a GFP-GCK-ALFA reporter and indicated candidate effector designs. Each point represents a transfection replicate and the dashed line at 1 indicates the mean of the negative-control effector (grey points). Effectors are ordered along the x-axis by mean normalized GFP:mCherry ratio. **B)** Scatterplot comparing measured effects from the singleton validation in HepG2 cells (x-axis) vs. HEK293T cells (y-axis). Pearson (r) and Spearman (ρ) correlation values are shown.

**Figure S6.**
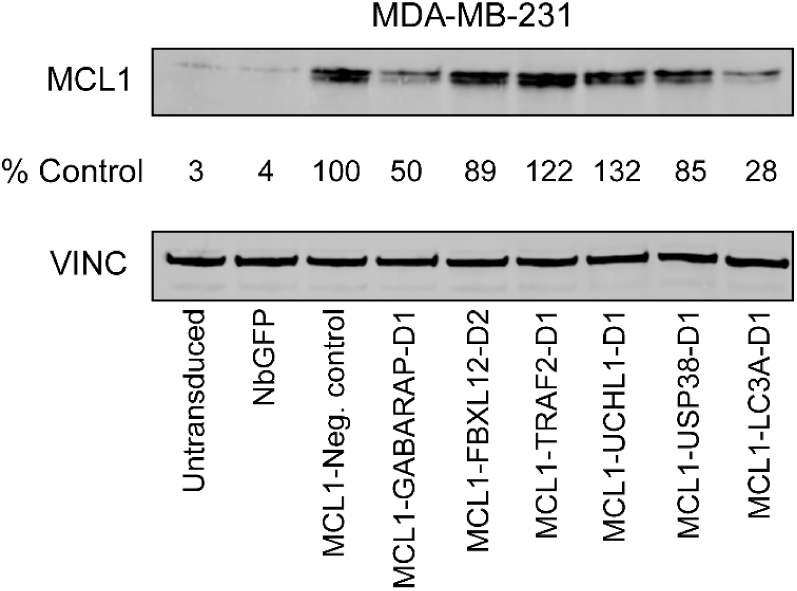
Assessment of effects of bifunctional MCL1-effector binder fusions on endogenous MCL1 levels. Western blot analysis of MCL1 levels in MDA-MB-231 cells, 72 hrs after transduction with indicated MCL1-effector binder fusions. Untransduced and GFP-nanobody (NbGFP) transduced cells serve as negative controls.

**Figure S7.**
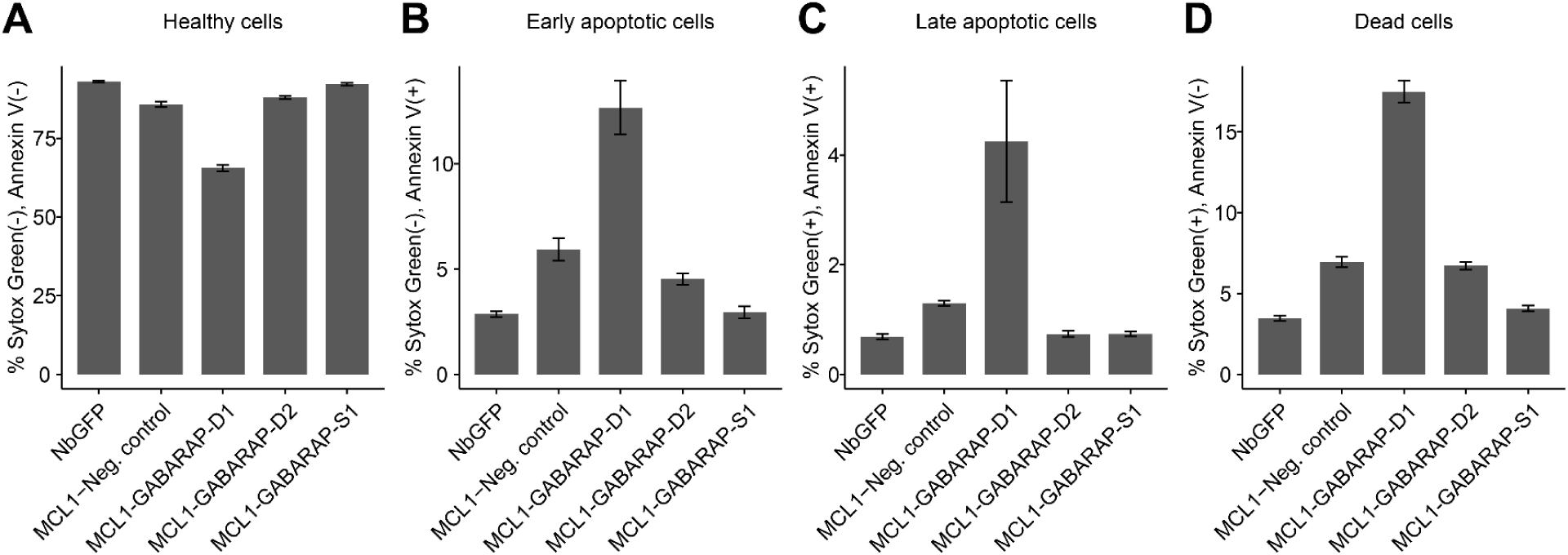
**Characterization of apoptosis induction by bifunctional MCL1-GABARAP binder fusions A-D**) Bar plots showing the percentage (y-axis) of cells in four states in the presence of various MCL1-effector binder fusions (x-axis). States were defined as: healthy (**A**), early apoptotic (**B**), late apoptotic (**C**), and dead (**D**) based on the indicated flow cytometry parameters (y-axis labels). Bars indicate the mean of three transduction replicates, error bars indicate ± SEM.

## SUPPLEMENTARY TABLES

**Table S1:** Oligos and plasmids used in this work

**Table S2:** Association table of barcode-effector handle pairs

**Table S3:** Barcode read counts from the LABEL-seq-screen

**Table S4:** Results of the LABEL-seq screen with replicate-level significance testing

**Table S5:** Results of the LABEL-seq screen with combined replicate significance testing

**Table S6:** Table of effector handle IDs and associated amino acid sequences

**Table S7:** Table of handle IDs and associated amino acid sequences used in validation experiments

## ACKNOWLEDGEMENTS

We thank members of the Fowler, Maly, Baker, and Shendure labs for helpful discussions on experimentation, data processing, analysis, and visualization. We disclose that language editing, proofreading and coding were supported by AI-based tools; these were not used for conceptual development or primary manuscript writing. The authors take full responsibility for the contents of this manuscript. C.C.S. was supported by the NIGMS (T32GM136534) and the NHLBI (F31HL168982). G.A. and Y.F. were supported by individual Jane Coffin Childs Postdoctoral Fellowships. G.A. and D.B. were supported by Gates Foundation grant INV-043758. This work was supported by the National Institutes of Health (R01HG010632 to J.S.), the Paul G. Allen Frontiers Group (Allen Discovery Center for Cell Lineage Tracing to J.S.), and the Brotman Baty Institute for Precision Medicine. D.B. and J.S. are investigators of the Howard Hughes Medical Institute.

## AUTHOR CONTRIBUTIONS

C.C.S. and G.A. conceptualized the application of *de novo* designed proteins for targeted protein degradation and stabilization with input from D.B. and J.S.. G.A. designed binders against FBXL12, GABARAP, GABARAPL2, MAP1LC3A, and UCHL1. S.S. designed binders against USP38 and TRAF2. C.C.S. designed and executed screening experiments with assistance from M.C. and input from J.J.S. and J.S.. C.C.S. analyzed screen data with input from J.S.. C.C.S. performed fluorescent reporter validation experiments with assistance from M.C. and D.S.L.. G.A. performed western blotting and apoptosis experiments with assistance from C.C.S.. Y.F. performed confocal microscopy of mitochondria with assistance from C.C.S.. C.C.S., G.A., D.B, and J.S. wrote the initial draft of the manuscript with input from all other authors. D.M.F., D.J.M., D.B., and J.S. supervised the project.

## DATA AVAILABILITY

Raw sequencing data and associated processed data are in the process of being uploaded to the Gene Expression Omnibus (GEO). The preprint will be updated to include the GEO identifier as soon as it is available. Additionally, the processed data files accompany this preprint (**Tables S2, S3**)

## MATERIALS & METHODS

### Cell lines and cell culture

HEK293T (CRL-3216), MDA-MB-231 (HTB-26), and HepG2 (HB-8065) cell lines were purchased from ATCC. circRNA-protein landing pad (RPLP) HEK293 cells were generated as described previously^2^. HEK293T, MDA-MB-231, HepG2, and HEK293 RPLP cells were cultured in DMEM (Gibco). All media were supplemented with 10% FBS (Hyclone) and 1% penicillin-streptomycin (Gibco). All cells were grown with 5% CO_2_ at 37 °C.

### Binder design

To design *de novo* binders, we generated 200 backbones using RFdiffusion and generated 10 sequences per backbone with ProteinMPNN^16^. We used AlphaFold2^27^ (AF2) with an initial guess and target templating to filter designs^33^. Design models that were predicted from AF2 with pae_interaction <20 were subjected to partial diffusion^15^ for backbone optimization, followed by two rounds of ProteinMPNN and AF2 filtering. Designs were selected by filtering on pae_interaction <8 and pLDDT >88.

### Binder library cloning

Barcoded binder libraries were cloned in a three step process (**Fig. S1B**). First, oligo libraries (Twist) encoding designed effector binders were PCR amplified and Gibson assembled (NEBuilder HiFi DNA Assembly, NEB) into XhoI/AgeI digested pCCS-266 plasmid backbone (**Fig. S1B, step I**). The Gibson assembly was cleaned (Clean and Concentrate kit, Zymo) and electroporated into *Escherichia coli* (NEB, C3020) which were then cultured at 30 °C for 20 hours at which point plasmid DNA was extracted (ZymoPURE II Plasmid Midiprep Kit). 1% of the electroporation was serially diluted and plated onto LB agar plates with ampicillin to quantify the number of transformants and ensure adequate library complexity.

Second, the library was barcoded via addition of a degenerate 16-mer DNA barcode to the backbone (**Fig. S1B, step II**). An oligonucleotide containing an internal stretch of 16 random bases was synthesized (IDT), PCR amplified to add Gibson homology handles, and Gibson assembled into BsrGI-digested backbone from **Step I**. As before, the Gibson assembly was cleaned and electroporated into *Escherichia coli*, and 1% of the electroporation was serially diluted and plated onto LB agar plates with ampicillin to quantify the number of transformants and ensure adequate library complexity. The barcode complexity was bottlenecked at this step by splitting various volumes of electroporated cells to different flasks (e.g. 1 mL of cells was split to 5 different flasks containing 25 µL, 75 µL, 150 µL, 200 µL, or 300µL) and these flasks were then cultured at 30 °C for 20 hours. The following morning, the number of transformants in the entire transformed volume was calculated and used to calculate the number of transformants inoculated into each flask. Plasmid DNA was extracted from the flask inoculated with the volume of transformed cells that most closely aligned to the target number of barcodes.

Finally, DNA encoding the ALFA nanobody (NbALFA) was cloned into the library (**Fig. S1B, step IV**). A gBlock containing the NbALFA coding sequence was synthesized (IDT), PCR amplified, and Gibson assembled into PacI/PmeI digested barcoded binder library from **Step II**. 1% of the electroporation was serially diluted and plated onto LB agar plates with ampicillin to quantify the number of transformants and ensure adequate library complexity.

### Binder-barcode subassembly

#### Preparation of subassembly library

Binders were associated to their corresponding barcodes via subassembly (Figure S1B, **step III**). Plasmid library DNA containing designed binders and degenerate 16-mer barcodes was subjected to two rounds of PCR. The first round of PCR utilized a forward primer that hybridizes upstream of the DNA barcode and a reverse primer that hybridizes downstream of the designed binder sequences. This first round PCR product was then subjected to a second round of PCR to introduce sequencing indices as well as P5 and P7 adapters for Illumina sequencing. The final amplicon was sequenced on a NextSeq2000 Illumina sequencer with Read 1 capturing the binder sequence and Read 2 capturing the barcode sequence.

#### Computational analysis of subassembly sequencing data

We implemented a Python script to pair each 16 nucleotide DNA barcode with its cognate designed binder sequence from demultiplexed, paired-end FASTQ files. The script requires as input a reference table of binder-encoding DNA sequences and their associated identifiers. From this reference, we first isolate the last 100 nucleotides of each binder sequence which is then reverse complemented. All possible single nucleotide changes of all reverse complemented sequences are then computed and stored for later reference.

Next, binder sequences are extracted from Read 1 by first finding a constant 10 nucleotide string immediately preceding the binder and subsequently extracting the next 100 nucleotides. These 100 nucleotides represent the last 100 nucleotides of the binder sequence and should correspond to a binder contained in the reverse complemented sequences processed previously. The extracted nucleotides are queried against the previously computed reference, and reads that match a single binder (or a single nucleotide variant of a single binder) are labelled with that binders corresponding identifier.

Barcodes are then extracted from Read 2 in a similar manner. Reads are parsed for a constant 10 nucleotide string immediately preceding the 16 nucleotide barcode. The subsequent 16 nucleotides representing the barcode sequence are then extracted.

Each paired read is written to a file containing a single barcode-binder pair per row. Reads for each barcode-binder pair are subsequently summed, and the distribution visualized as a histogram (**Fig. S1C**). From the histogram, we set a threshold (i.e. 10 reads per barcode-binder pair) above which we consider barcode-binders pairs to be true. As a second filter, we compute the proportion of all unique associations for each barcode (e.g. 80% of barcode ATGC reads map to binder 1, 15% to binder 2, etc…). We take the maximum of the proportion value for each barcode and visualize it as a histogram (**Fig. S1D**) which shows that the vast majority of barcodes have a high proportion of their reads mapped to an individual binder. Barcodes with a maximum read proportion ≥ 80% are retained. These barcode-binder pairs (i.e. those with ≥ 10 reads & ≥ 80% of reads associated with a single binder) are then saved as a lookup table. <u>LABEL-seq screen</u>

#### Library recombination

Barcoded binder libraries were recombined into RPLP HEK293T cells at a genomically integrated landing pad as previously described^2^. On day 0, four 15 cm tissue culture dishes were individually seeded with 18×10^6^ RPLP HEK293T cells in complete DMEM. On day 1, cells were transfected with 20 µg of barcoded binder library (**Fig. 1C**) and 3 µg pCAG-NLS-HA-Bxb1 (Addgene #51271) using Lipofectamine 3000 according to the manufacturers instructions, and the media was replaced with complete DMEM on day 2. Expression of genomically integrated binders was induced on day 3 by addition of doxycycline (Sigma-Aldrich #D9891) to a final concentration of 2 µg mL^-1^. Recombined cells were then selected on day 5 by treating cells with 1 nM AP-1903 (Selleckchem S9726) for 6 hours, after which media was replaced with complete DMEM supplemented with 2 µg mL^-1^ doxycycline. On day 7, 2 µg mL^-1^ puromycin (Thermo A1113803) was added to the media to maintain selection for recombined cells and cells were expanded until day 13 at which point cells were trypsinized, pelleted, resuspended in CELLBANKER 1 (Amsbio 11884) and frozen at −80 °C.

#### Immunoprecipitation of reporter proteins and their associated circular RNA barcodes

Frozen aliquots of RPLP HEK293T recombined with the barcoded binder library were thawed and expanded in complete DMEM supplemented with 2 µg mL^-1^ doxycycline. Six 15 cm tissue culture dishes were individually seeded with 20×10^6^ recombined RPLP HEK293T cells expressing the barcoded binder library.

These cells were transfected with 30 µg of dual POI/Standard RBP expression plasmid 24 hours post-seeding using Lipofectamine 3000 according to the manufacturer’s instructions. Cells were harvested and reporter proteins immunoprecipitated 48 hours post-transfection.

A 5x concentrated modified RIPA buffer (5xModRIPA; 250 mM Tris, 750 mM NaCl, 20mM MgCl_2_, and 25% glycerol) was prepared before immunoprecipitation and stored at 4 °C. The following buffers were prepared fresh on the day of immunoprecipitation:

Resuspension buffer: 1xModRIPA, 2 mM PMSF (Millipore Sigma 10837091001), 1x protease inhibitor cocktail (Thermo 78430), 20 units mL^-1^ SUPERaseIn RNase Inhibitor (Thermo AM2696), 300 nM synthetic decoy RNA.

Lysis buffer: 1xModRIPA, 1x protease inhibitor cocktail (Thermo 78430), 20 units mL^-1^ SUPERaseIn RNase Inhibitor (Thermo AM2696), 0.2% Igepal CA-630 (Sigma-Aldrich I8896-50ML).

Wash buffer: 1xModRIPA, 0.1% Igepal CA-630 (Sigma-Aldrich I8896-50ML).

The six 15 cm tissue culture plates were trypsinized and combined into two replicates each consisting of cells from 3 individual plates. Replicate pools were counted and 9×10^7^ cells per replicate were pelleted by centrifugation at 500×g for 5 minutes after which supernatant was removed and the samples transferred to ice. Pellets were resuspended in 4.5 mL of ice cold resuspension buffer and then lysed by addition of 4.5 mL of lysis buffer followed by a 10 minute incubation on ice. Cell lysates were then transferred to 2 mL tubes and insoluble material cleared by centrifugation at 17,000×g for 10 minutes at 4 °C.

For each replicate, 300 µL of Pierce anti-c-Myc magnetic beads (Thermo 88842) and 300 µL of anti-FLAG M2 magnetic beads (Sigma M8823) were prepared by collecting beads on a magnet, removing supernatant, and resuspending with 1.5 mL (i.e. 5x bead volume) of wash buffer. The bead & wash buffer solution was removed from the magnet, inverted to mix, and then washed an additional 3 times with a 5x bead volume of wash buffer. After the final wash, beads were resuspended in 300 µL of wash buffer.

Lysates from each replicate were split between FLAG (3 mL lysate) and Myc (6 mL lysate) immunoprecipitations, and 300 µL of washed beads were added to the corresponding lysate aliquot. Samples were then rotated end-over-end at 4 °C for 3 hours in a cold room, after which beads were collected on a magnet, washed 3 times with 3 mL of wash buffer (i.e. 10x bead volume). After the final wash, beads were collected on a magnet, supernatant was removed, and beads were resuspended in 150 µL of PBS followed by 450 µL of TRIzol LS reagent (Invitrogen 10296010). Resuspended beads were then frozen at −20 °C.

#### RNA isolation and generation of DNA sequencing library

Frozen beads were thawed on ice, beads collected on a magnet, and supernatant was transferred to a clean tube. Beads were washed with an additional 200 µL of TRIzol LS prediluted 1:4 with PBS (e.g. 250 uL of PBS + 750 uL Trizol-LS) and that 200 µL was transferred to the same tube. RNA was isolated from the supernatant using a Direct-zol RNA Miniprep (Zymo R2050) according to the manufacturer’s instructions. RNA was eluted in 50 µL of nuclease-free water. 45 µL of eluted RNA was mixed with 2 µL of TURBO DNase (Invitrogen AM2238), 5 µL of 10X Turbo DNase buffer, and 3 µL of nuclease free water and incubated at 37 °C for 30 minutes. RNA was cleaned using a Monarch Spin RNA Cleanup Kit (NEB T2040S) and eluted in 25 uL of nuclease free water.

The entire volume of eluted RNA was then reverse transcribed into cDNA using SuperScript IV Reverse Transcriptase (Invitrogen 18090010). First, 25 µL of cleaned RNA was mixed with 10 uL 1 uM oCCS-1267 reverse transcription primer, 5 uL 10 mM dNTPs, and 25 uL nuclease free water. This mixture was incubated at 65 °C for 5 minutes, and then transferred to ice. Reverse transcriptase reaction mix consisting of 20 µL of 5x SSIV buffer, 5 µL 100 mM DTT, 5 µL nuclease free water, and 5 µL of SuperScript IV Reverse Transcriptase were combined together and then mixed with 65 µL of previously annealed RNA and RT primer. Samples were incubated at 55 °C for 30 minutes, followed by an incubation at 80 °C for 10 minutes and then at 4°C.

To convert the cDNA into an Illumina sequencing library, the entire 100 µL volume of cDNA was spread across eight 50 µL PCR reactions each consisting of 0.25 uL 100 µM oCCS-1266 forward primer, 0.25 µL 100 µM oCCS-1268 reverse primer, 12.5 µL of RT reaction product, 0.25 µL of 100X SYBR Green (ThermoFisher S7567), 25 µL of KAPA HiFi HotStart ReadyMix (Roche KK2602), and 11.75 µL of nuclease free water. Cycling parameters were 3 minutes at 95 °C followed by 15 cycles of 20 seconds at 98 °C, 15 seconds at 75 °C and 20 seconds at 72 °C. Reactions were monitored on a realtime qPCR machine (CFX Opus, Bio-Rad) and terminated while the fluorescence signal was in the log phase. The 8 PCR reactions for each sample were pooled, cleaned using 1.5x AMPure XP beads (Beckman Coulter Life Sciences A63882), and eluted in 240 µL of Buffer EB (Qiagen 19086). Illumina sequencing adapters and indices were added to each sample in a second PCR reaction consisting of 20 μL 1st round PCR product, 100 μL of 2x NEBNext master mix, 10 μL 10 μM P5 indexing primer (for example oCCS-P5-1), 10 μL of 10 µM P7 indexing primer (for example oCCS-P7-1), and 60 μL of nuclease-free water and cycling parameters of 30 seconds at 98 °C followed by 5 cycles of 10 seconds at 98 °C, 30 seconds at 63 °C and 15 seconds at 72 °C.

Reverse transcription of a circular RNA produces concatameric cDNA products which result in laddered PCR amplicons^34^. Indexed PCR reactions were run on a 6% polyacrylamide gel (Invitrogen EC62652BOX) at 200V for 35 minutes, after which the gel was stained with SYBR Gold (Invitrogen S11494). Bands were visualized on a light box and the smallest band in the laddered product was excised from the gel. DNA was purified from the gel following a previously published protocol^35^.

Purified products were quantified on an Agilent 4200 TapeStation, pooled, and sequenced on a Illumina NextSeq 2000 sequencer.

#### Computational analysis of barcodes

Barcode-containing sequencing reads were processed with custom Python scripts to extract barcodes and assign corresponding binder identities. Demultiplexed paired-end FASTQ files were then processed as follows. The barcode sequence was extracted from Read 1 as the nucleotide stretch between predefined flanking sequences, followed by computation of its reverse complement. This barcode was then queried against the lookup table of barcode-binder pairs (see “Computational analysis of subassembly sequencing data” section) to retrieve the corresponding binder identifier. The sample name, read identifier, extracted barcode, extracted barcode reverse complement, binder assignment, and mean Phred quality across the entire read were recorded. The dataset was written to a CSV file containing a single read and its associated information per row.

Downstream analysis of the binder–barcode sequencing output was performed in R. The CSV file containing read-level annotations was imported, and filtered to 1) retain reverse-complement barcodes of the expected length, 2) exclude barcodes lacking an associated binder, and 3) exclude . Remaining reads were collapsed to unique combinations of barcode, binder ID, pulldown (i.e. FLAG or Myc), and replicate with an associated read count.

To compare results between replicates, per-sample read proportions were calculated from summed read counts of each barcode-binder pair in each pulldown in each replicate. Barcodes were retained only if they were detected at a non-negligible proportion in both replicates (per-barcode proportion ≥10^−7^ in both replicate 1 and replicate 2). Only binders with ≥ 3 barcodes in each replicate were retained and a small pseudocount (10^−6^) was added to each read proportion. Within each replicate, differences in the FLAG and Myc read proportions were tested by a paired Wilcoxon signed-rank test utilizing all barcodes belonging to the same binder. P values were corrected for multiple-hypothesis testing using the Benjamini–Hochberg method.

We then pooled all read proportions and recomputed significance in order to increase our statistical power. Similar to the between-replicate test, barcodes with a frequency ≥10^−7^ in both replicates were retained and binders with ≥ 3 barcodes in each replicate were retained and a small pseudocount (10^−6^) was added to each read proportion. Differences in the FLAG and Myc read proportions were again tested by a paired Wilcoxon signed-rank test utilizing all barcodes belonging to the same binder and P values corrected for multiple-hypothesis testing using the Benjamini–Hochberg method.

### Singleton validation of screen effects with a fluorescent protein abundance reporter

A stable, polyclonal HEK293T cell line harboring an EF1a-driven reverse tetracycline transactivator (rtTA) and a doxycycline-inducible TRE-3xFLAG-GFP-GCK-ALFA-IRES-mCherry plasmid was generated by piggyBac transposition. A negative control line was harboring an EF1a-driven rtTA and a reporter in which the ALFA tag was deleted (i.e. 3xFLAG-GFP-GCK-IRES-mCherry) was generated in parallel. 2×10^5^ of these cells were seeded in wells of a 24 well plate and transfected 24 hours later with 500 ng of a plasmid encoding a NbALFA-binder fusion and mTagBFP2 driven by CMV and EF1a promoters respectively. Reporter expression was induced 24 hours post transfection via addition of 100 ng/mL doxycycline to the culture media. 72 hours post-transfection (48 hours post-induction) cells were trypsinized, pelleted at 500×g for 5 minutes, resuspended in FACS buffer (PBS supplemented with 10% FBS), and analyzed on a Attune Flow Cytometer (Thermo). These results are presented in **Figure 3A-3B**.

Binder-induced changes in the GFP:mCherry ratio was also measured in HepG2 cells with several changes. Here, HepG2 cells were seeded (1.1×10^5^ cells per well of a 24 well plate) and co-transfected with 500 ng each of constitutive binder and reporter plasmids (i.e. CMV-NbALFA-binder-EF1a-mTagBFP and CMV-3xFLAG-GFP-GCK-IRES-mCherry, respectively).

### Analysis of fluorescent protein abundance reporter flow cytometry data

Flow cytometry experiments were analyzed using FlowJo. Cells were gated from all events (forward scatter area versus side scatter area) followed by gating on single cells (forward scatter area versus forward scatter height). From the population of single cells, mCherry positive cells were gated, followed by mTagBFP2 positive cells. A GFP:mCherry ratio parameter was derived and applied to this population.

### Assessment of endogenous MCL1 protein degradation by Western blotting

To prepare lentiviral particles encoding MCL1-degrader binder fusions, 1.2×10^6^ HEK293T cells were seeded in individual wells of a 6 well plate one day before transfection with pMDLg/pRRE (Addgene #12251), pRSV-Rev (Addgene #12253), pMD2G (Addgene #12259) and 750 ng of binder-encoding lentiviral vector. Transfection media was changed 24 hours post transfection. 48 hours post-transfection, virus containing supernatant was collected, centrifuged at 500×g for 5 minutes, and filtered through a 0.45 µm filter.

3×10^5^ MDA-MB-231 cells were seeded in 2 mL of complete DMEM in individual wells of a 6 well plate 24 hours before transduction. To transduce cells, 1 mL of DMEM was removed from each well and replaced with 1 mL of freshly prepared and filtered lentiviral supernatant. Media were replaced with 2 mL of complete DMEM 24 hours post-transduction. 72 hours post-transduction, cells were washed 3 times with DPBS and lysed with RIPA Lysis Buffer supplemented with protease inhibitor, phosphatase inhibitor, and benzonase on ice for 30 minutes. Cells were scraped and spun down at 21,000×g for 15 minutes at 4 °C. The supernatant was collected and the concentrations were normalized through a BCA assay. 45 μg of lysates were loaded onto 4-12% Bis-Tris Gel and separated by SDS-PAGE. The gel was then transferred to a nitrocellulose membrane, and blocked with Intercept Blocking Buffer (LICOR) for 1 hour at room temperature. The blots were stained with primary antibody overnight at 4 °C, washed 3 times with TBS-T, and stained with secondary antibody for 1 hour at room temperature. A LICOR Odyssey CLx Imager was used to image and quantify the blot.

### Apoptosis Assay

1.1×10^5^ MDA-MB-231 cells were seeded in 0.3 mL of complete DMEM in individual wells of a 48 well plate 24 hours before transduction. The following day, 150 µL of DMEM was removed from each well and replaced with 150 µL of freshly prepared lentiviral supernatant. Media was replaced with 300 µL of complete DMEM 24 hours post-transduction. 72 hours post-transduction, cells were trypsinized, quenched with media, and moved to a 96-well-U-bottom plate. Cells were spun down and washed once with DPBS. Each well was incubated with 50 µl Annexin Buffer (10 mM HEPES, 140 mM NaCl, and 2.5 mM CaCl2, pH 7.4) and 2.5 µl of Annexin V AF647 Conjugate (Thermo Fischer A23204) for 15 minutes at rt. Afterwards, 150 µl of Annexin buffer with Sytox Green was added, and cells were analyzed by Attune Flow Cytometer.

### Confocal microscopy

3×10^4^ HEK293T cells were seeded in 96-well glass bottom plate (Cellvis #P96-1.5H-N) and transfected 24 hours later with 25 ng of ALFA-GFP-FIS1-IRES-mCherry and 75 ng of NbALFA-degrader-EF1a-mTagBFP plasmids. Media was replaced 24 hours post-transfection with complete DMEM supplemented with 125 nM MitoTracker Deep Red FM dye (ThermoFisher Scientific #M22426) and cells were incubated in a tissue culture incubator at 37°C. After staining for 30 minutes, media was replaced with complete DMEM and confocal imaging was performed using a spinning-disk confocal microscope. As described previously ^36^, a Yokogawa CSU-W1 SoRa spinning disc confocal device is attached to a Nikon ECLIPSE Ti2 microscope. Excitation light was emitted from lasers housed inside of a Nikon LUNF 405/488/561/640NM 1F commercial launch. Emission light was directed by a quadband dichroic mirror (Semrock, Di01-T405/488/568/647-13×15×0.5) and filtered by one of four single-bandpass filters (DAPI, Chroma, ET455/50M; ATTO 488, Chroma, ET525/36M; ATTO 565, Chroma, ET605/50M; Alexa Fluor 647, Chroma, ET705/72M) and focused onto an Andor Sona 4.2B-11 camera. Specific settings for this work were as follows: Excitation light was emitted for 405 nm, 488 nm, 561 nm, or 640 nm lasers at 30%, 35%, 30%, and 35% of maximal intensity, respectively. The exposure times of 600, 200, 200, and 200 ms were used for 405 nm, 488 nm, 561 nm, and 640 nm, respectively. A 100x N.A. 1.49 Apo oil immersion objective lens and a 2.8x lens in the SoRA unit were used for super-resolution imaging, resulting in an effective pixel size of 39.3 nm. Z-stacks were captured for all channels at a 0.3 μm step size for 51 steps using the “Z then channel” sequence. Nikon Elements AR 5.20.00 was used to acquire microscopy images on the Nikon Ti2 system. Images were processed in ImageJ (Fiji).

